# Genomic inbreeding trends in the global Thoroughbred horse population driven by influential sire lines and selection for exercise trait-related genes

**DOI:** 10.1101/770545

**Authors:** Beatrice A. McGivney, Haige Han, Leanne R. Corduff, Lisa M Katz, Teruaki Tozaki, David E. MacHugh, Emmeline W. Hill

## Abstract

The Thoroughbred horse is a highly valued domestic animal population under strong selection for athletic phenotypes. Here we present a high resolution genomics-based analysis of inbreeding in the population that may form the basis for evidence-based discussion amid concerns in the breeding industry over the increasing use of small numbers of popular sire lines, which may accelerate a loss of genetic diversity. In the most comprehensive globally representative sample of Thoroughbreds to-date (*n* = 10,118), including prominent stallions (*n* = 305) from the major bloodstock regions of the world, we show using pan-genomic SNP genotypes that there has been a highly significant decline in global genetic diversity during the last five decades (*F*_*IS*_ *R*^2^ = 0.942, *P* = 2.19 × 10^−13^; *F*_*ROH*_ *R*^2^ = 0.88, *P* = 1.81 × 10^−10^) that has likely been influenced by the use of popular sire lines. Estimates of effective population size in the global and regional populations indicate that there is some level of regional variation that may be exploited to improve global genetic diversity. Inbreeding is often a consequence of selection, which in managed animal populations tends to be driven by preferences for cultural, aesthetic or economically advantageous phenotypes. Using a composite selection signals approach, we show that centuries of selection for favourable athletic traits among Thoroughbreds acts on genes with functions in behaviour, musculoskeletal conformation and metabolism. As well as classical selective sweeps at core loci, polygenic adaptation for functional modalities in cardiovascular signalling, organismal growth and development, cellular stress and injury, metabolic pathways and neurotransmitters and other nervous system signalling has shaped the Thoroughbred athletic phenotype. Our results demonstrate that genomics-based approaches to identify genetic outcrosses will add valuable objectivity to augment traditional methods of stallion selection and that genomics-based methods will be beneficial to actively monitor the population to address the marked inbreeding trend.

**Author Summary:** In the highly valuable global Thoroughbred horse industry, there is no systematic industry-mediated genetic population management. Purposeful inbreeding is common practice and there is an increasing use of popular sires. Inbreeding can lead to population health and fertility decline, but there is little objective genomics-based data for the Thoroughbred to catalyse action and support changes in breeding practices. Here, we describe the most comprehensive genetic analysis in the population among 10,000 Thoroughbreds from the major bloodstock regions of the world and reveal a highly significant increase in inbreeding during the last five decades. The main drivers of genetic diversity are the most influential ‘breed-shaping’ sire lines, *Sadler’s Wells, Danehill* and *A.P. Indy*. We identified genomic regions subject to positive selection containing genes for athletic traits. Our results highlight the need for population-wide efforts to proactively avert the potential increase of deleterious alleles that may impact on animal health in order to safeguard the future of a breed that is admired for its athleticism and enjoyed for sport worldwide.

## Introduction

The Thoroughbred is among the fastest animals selected by humans for sport, originating from *“the commingled blood of Arabs, Turks and Barbs”* [1] crossed with local British and Irish mares [2]*“but selection and training have together made him a very different animal from his parent-stocks”* [1]. The Thoroughbred is now a large (N ~500,000) global breed but, in the context of modern horse breeds, it has very low genetic diversity [3, 4] due to the limited foundation alleles at the establishment of the stud book [5–7] and restriction of external gene flow subsequent to the closing of the population. In Thoroughbred horse breeding selection of potential champion racehorses is a global multi-billion-dollar business, but there is no systematic industry-mediated genomic selection or genetic population management. We hypothesised that the market-driven emphasis on highly valuable pedigrees and the common practice of inbreeding to successful ancestors in attempts to reinforce favourable variants in offspring has resulted in a global reduction in genetic diversity. Here, we apply population genetics approaches to assess temporal population-wide variation across the longest time span to-date for this population, and for the first time perform a comparative analysis of genetic diversity among the major bloodstock regions of the world. Additionally, we identify genes of interest for the Thoroughbred phenotype in signatures of positive selection that are likely to be most impacted by inbreeding.

## Results and Discussion

### Genetic diversity driven by ‘breed-shaping’ stallions within a highly homogeneous global population

We evaluated genetic diversity in a principal component analysis (PCA) of the genetic relatedness matrix between Thoroughbreds and representatives of putative founding populations and within the global Thoroughbred population. We show that the Thoroughbred breed is divergent from the founding populations (S1-S2 Figures and S1 Text). However, although the population is geographically dispersed, with the majority of horses located in Australasia (ANZ), Europe (EUR), Japan (JAP), North America (NAM) and South Africa (SAF), the Thoroughbred (*n* = 10,118, b.1970-2017) is largely genetically homogeneous maintained in a single cluster, *albeit* with some level of geographic population structure particularly towards EUR samples. The outliers driving diversity from the main cluster are prominent stallions (S3-S4 Figures) and visualisation of relatedness among the stallion population (*n* = 305) reveals partitioning in PC1 and PC2 apparently driven by the ‘breed-shaping’ sire lines in each region; *Sadler’s Wells* (b. 1981 NAM) in EUR; *Danehill* (b. 1986 NAM) in ANZ and *A.P. Indy* (b. 1989 NAM) in NAM (Figure 1, S5 Figure). PCA plots illustrating the genetic diversity in each of the major regions EUR, ANZ and NAM are provided in S6-S11 Figures.

**Figure 1:**
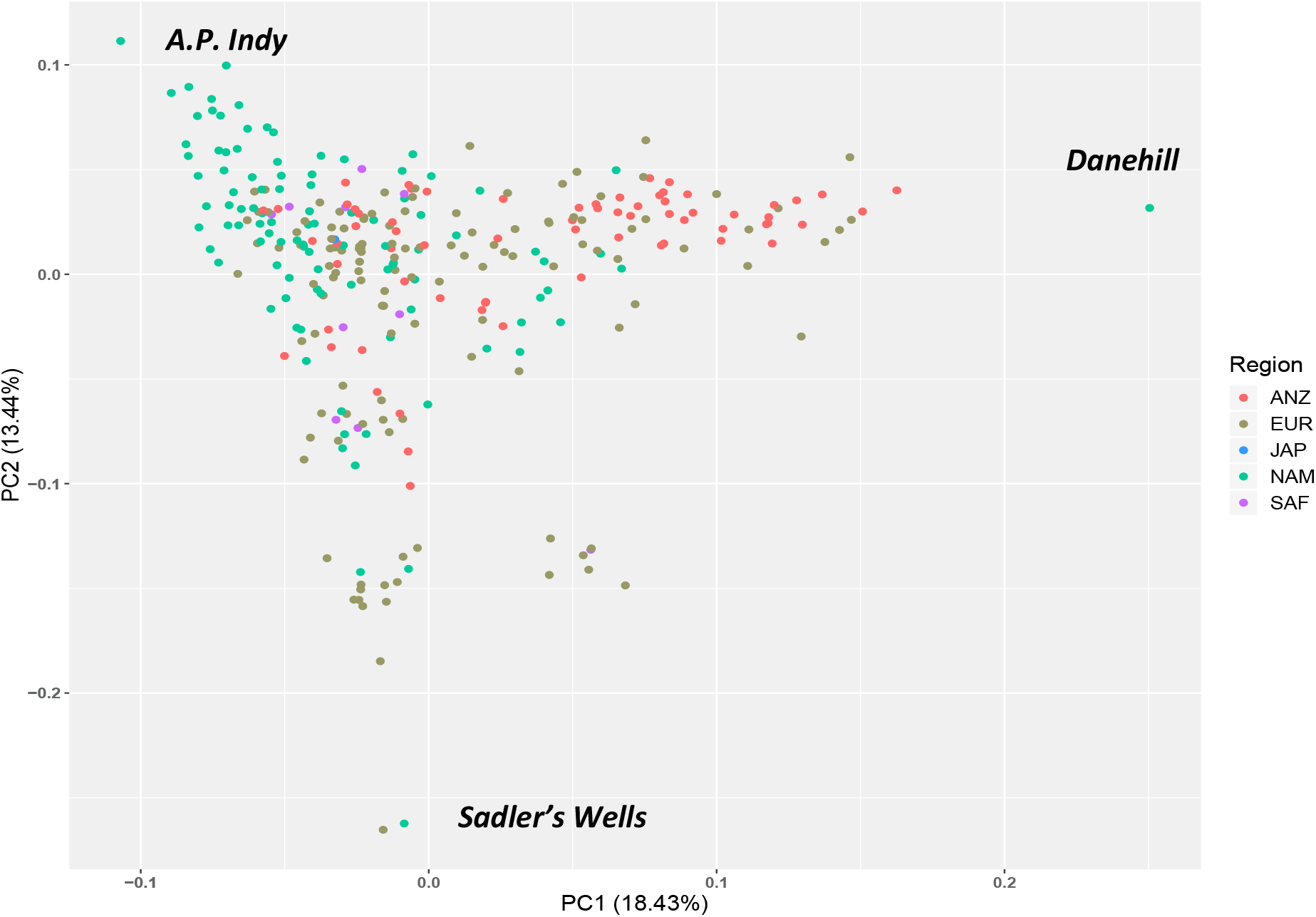
Principal component analysis plot of the genetic relatedness matrix based on genotype data for prominent global Thoroughbred stallions (*n* = 305). Individuals are colour coded based on region of birth: Australia / New Zealand (ANZ), red; Europe (EUR), green; Japan (JAP), blue; North America (NAM), light green; South Africa (SAF), purple.

### A marked increase in inbreeding in the Thoroughbred population over five decades

Here, in the most comprehensive genetic analysis performed in this population (*n* = 10,118) we show a striking temporal increase in inbreeding and regional variance across the global Thoroughbred population during the last five decades. Individual inbreeding coefficients (*F*_IS_) estimated using 9,212 pruned SNPs and runs of homozygosity (*F*_ROH_) (minimum length 1 Mb) characterised using an unpruned set of 46,478 SNPs revealed a highly significant increase in inbreeding over time (*F*_IS_: *R*^2^ = 0.942, *P* = 2.19 × 10^−13^; *F*_ROH_: *R*^2^ = 0.88, *P* = 1.81 × 10^−10^) (Figure 2) with the greatest rate of change observed since the 2000s (S12 Figure). A linear regression model was used to test for significance and directionality of change in annual mean inbreeding with respect to year of birth (S12 Figure). Similar trends were observed within each geographic region (S13-S16 Figures). Results from Student’s t-tests indicated that inbreeding estimates were higher in EUR compared to ANZ (*F*_IS_, *P* =1.9 × 10^−21^; *F*_ROH_, *P* = 0.043) and NAM (*F*_IS_, *P* < 3.1 × 10^−19^; *F*_ROH_, *P* = 0.00059). A temporal reduction of Thoroughbred genetic diversity has previously been reported among much smaller sample sets from single geographic regions [8, 9] Expansion of the timeline to include the most recent decade using a sample size more than 20-fold larger indicates that despite industry cognizance of inbreeding and previous cautions [8], there has been no arrest in the rate of increase in inbreeding and it is a global, population-wide phenomenon.

**Figure 2:**
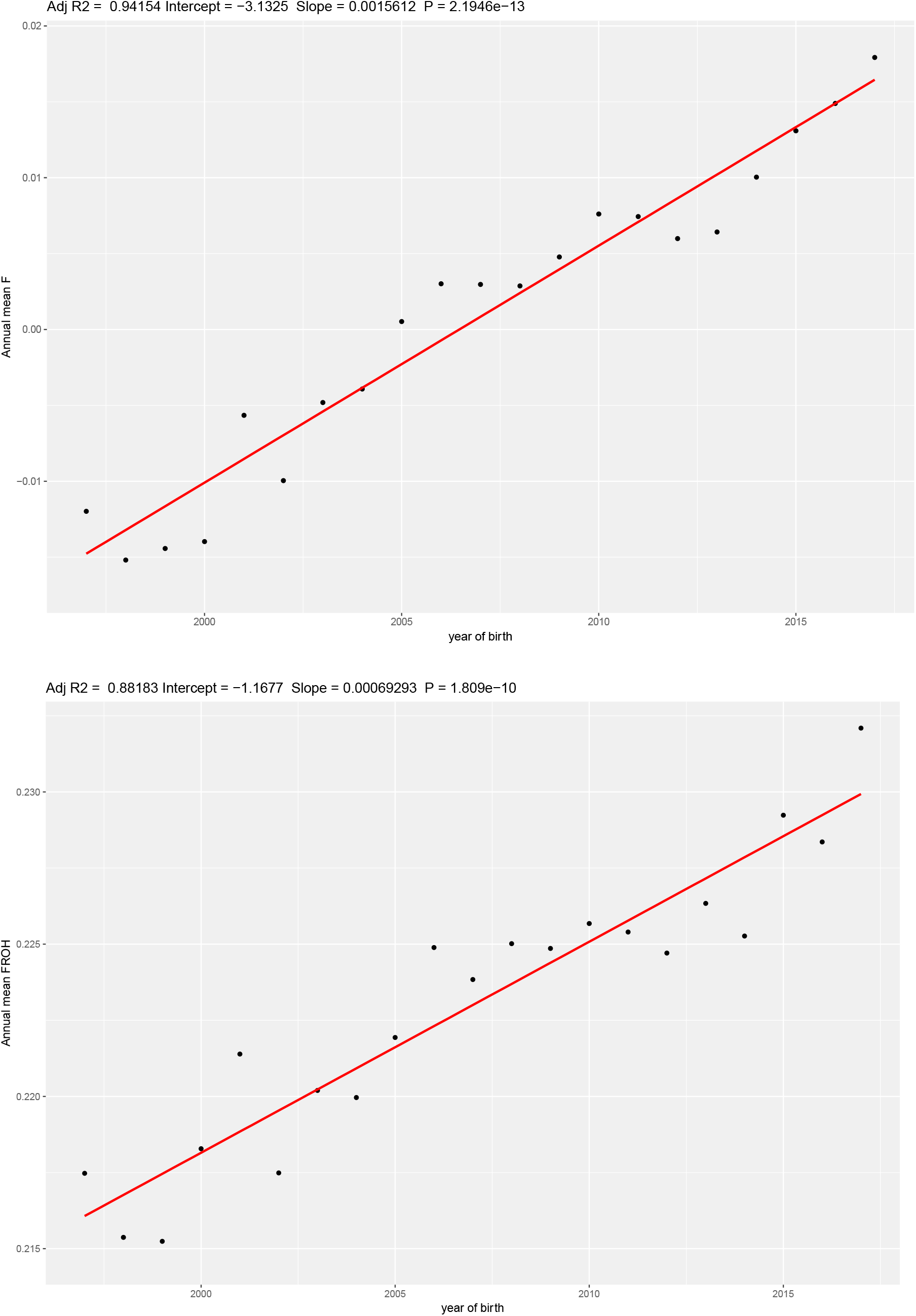
The regression of the mean annual *F*_IS_ [top] and *F*_ROH_ [bottom] on year of birth for *n* = 10,118 Thoroughbred horses born between 1996 and 2017.

Breeding practices that promote inbreeding have not resulted in a population of faster horses [4, 10] and our results, generated for the first time using a large cohort of globally representative genotypes, corroborate this. When evaluating Timeform handicap ratings for EUR horses, no association was found between *F*_IS_ and racing performance for horses born between 1996-2013 (*n* = 1,886, *r*^2^ = 0.034, *P* = 0.780) and 2009-2013 (*n* = 1,065, *r*^2^ = −0.014, *P* = 0.65). Purposeful inbreeding attempts to duplicate favourable gene variants that are selected at each successive generation, but homozygosity arising from inbreeding tends to be associated with decreased trait values since the proportionate trait gain arising from beneficial mutations is generally limited by pleiotropy [11]. Also, among domesticates, there is an increased mutational load and a higher proportion of deleterious alleles in regions under selection [12–15]. Therefore, there may be a genetic cost of inbreeding, which in dogs has negatively influenced intra-breed genetic diversity and increased the frequency of disease-causing alleles [15, 16] and in cattle, is associated with decreases in milk, fat and protein yields and negatively impacts reproductive traits [17].

While inbreeding depression generally negatively impacts health and fertility through the accumulation of deleterious alleles, the ‘tipping point’ at which there is an irreversible accumulation of unfavourable mutations is not currently known. In horses mutational load and purging of deleterious alleles has been assessed in whole genome sequence data and revealed a significant relationship between inbreeding and mutational load [18]. Surprisingly, Thoroughbreds appear to have a lower than expected mutational load, suggested to be due to effective purging through negative selection on phenotypes. This may be facilitated in the Thoroughbred in practice by the unusually large census population size relative to the effective population size, with a high proportion of horses that do not ever race [19]. However, the results from that study are likely not representative of the genetics of the current breeding population in the major bloodstock regions of the world; the sample cohort of Thoroughbreds examined was small (*n* = 19) and all the horses were registered Thoroughbreds in Korea. For example, ROH identified in that study were up to 11 Mb, whereas we have detected here ROH up to 40 Mb (S1 Table); increasing ROH increases the likelihood of deleterious alleles being exposed.

It is interesting to consider also that many of the performance limiting genetic diseases in the Thoroughbred do not generally negatively impact on suitability for breeding; some diseases, with known heritable components, are successfully managed by surgery (osteochondrosis desicans, recurrent laryngeal neuropathy), nutritional and exercise management (recurrent exertional rhabdomyolysis) and medication (exercise induced pulmonary haemorrhage). This facilitates retention of risk alleles in the population and enhances the potential for rapid proliferation of risk alleles if they are carried by successful stallions. Furthermore, since single gene variants have not yet been identified for these diseases, it is likely that they result from polygenic additive genetic variation that may not be easily exposed and purged by negative selection on individual ROH. In order to fully understand the consequences of inbreeding in the Thoroughbred, inbreeding measures should be regressed on large population-scale phenotypes and the extent of mutational load should be determined in a large cohort of representative samples.

There are currently concerns about the impact of the increasing use of small numbers of stallions in the breeding population on inbreeding and population viability. While the census population size of the Thoroughbred is large, a more appropriate method to assess population health is to estimate the effective population size (*N*_e_), an estimate of the size of an idealised population that is representative of the genetic diversity in the actual population [20–22]. To better understand the impact of inbreeding on diversity in the population we estimated *N*_e_ for the global population and regional populations using the subset of horses born 2013-2017 (*n* = 3,341) to represent the current breeding population. The global population had *N*_e_ = 330 and among regional cohorts *N*_e_ was highest in NAM (*N*_e_ = 226) and lowest in SAF (*N*_e_ = 93); also ANZ (*N*_e_ = 197), EUR (*N*_e_ = 198). The observation that *N*_e_ was higher when all regions were considered together indicates that there is opportunity to exploit regional variation to improve genetic diversity in the global population. This may be achieved in practice, for example, by selecting mares in the breeding population that are genetically distant from the stallion, or more broadly by the permanent movement of stallions to regions that have a genetically diverse population of mares. A comparison of *N*_e_ with census population sizes (*N*_c_, included here as *N*_c_ = 100,000 for NAM, EUR, ANZ and SAF and *N*_c_ = 500,000 global) indicates that the genetic variation in the population deviates substantially from expectations. *N*_e_/*N*_c_ was <0.2% in all regional cohorts and was <0.1% when the population was considered as a whole. In most wild species *N*_e_ is 10-50% of the census population size and among domesticates *N*_e_/*N*_c_ has a median of 3% [23]. A limitation to interpretation of the results from this study is the potential bias in the samples used, many of which came from the same breeding farms. Also, in some cases, the sample sizes are relatively small (e.g. SAF) and may not be representative of the entire population. Nonetheless, these results highlight the need for a systematic evaluation of genetic diversity that may be applied for longitudinal monitoring of the Thoroughbred population.

### Selection for the Thoroughbred phenotype

Inbreeding is a consequence of selection for favourable traits. Here, to systematically identify genes that have been targets of selection for the Thoroughbred athletic phenotype, and may be most impacted by inbreeding, we used the composite selection signal (CSS) approach, a weighted measure that combines the signals from identification of highly differentiated loci (*F*_ST_), increase in selected allele frequency (*ΔSAF*) and cross-population extended haplotype homozygosity (*XP-EHH*) tests [24, 25], to analyse 48,896 pan-genomic SNPs genotyped in elite Thoroughbreds (*n* = 110) and non-Thoroughbreds (*n* = 84) representing putative founder populations. We identified 15 significant candidate selected genomic regions (S2 Table, S17 Figure), defined as clusters of ≥5 SNPs among the top 1% of the smoothed CSS statistic result (−log_10_*P*), seven of which overlapped with previously reported genomic regions with evidence for selection in Thoroughbreds [26, 27] (Table 1). As a high prevalence of runs of homozygosity (ROH) can inform on selection, the top 1,000 SNPs ranked by the percentage of individuals with a certain SNP located within a ROH were extracted for ROH >1Mb and >5Mb (S3-S4 Tables, S18 Figure). Among the CSS regions, there was substantial overlap with regions with a high prevalence of ROH, particularly shorter ROH suggesting recent/ongoing selection.

**Table 1:**
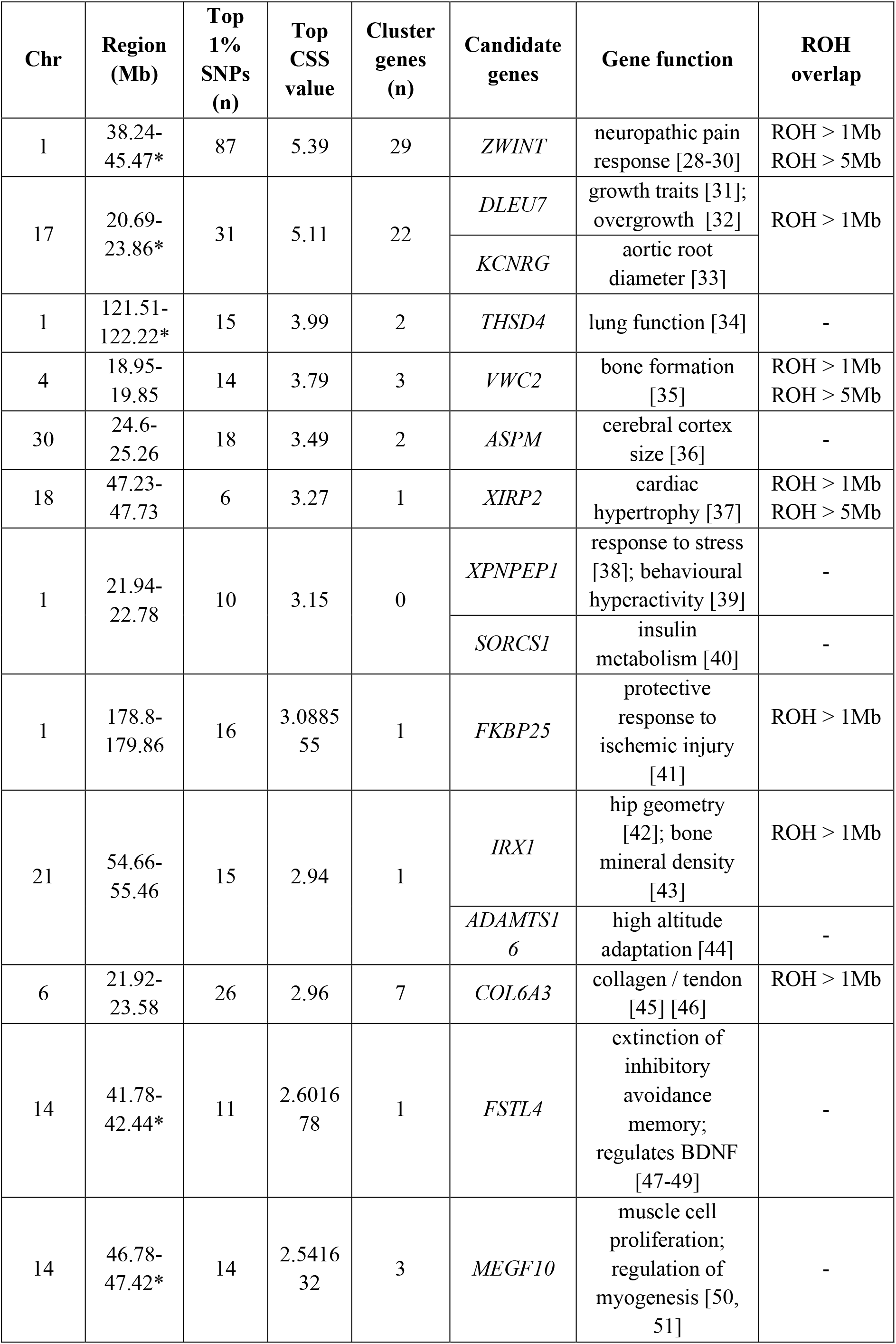
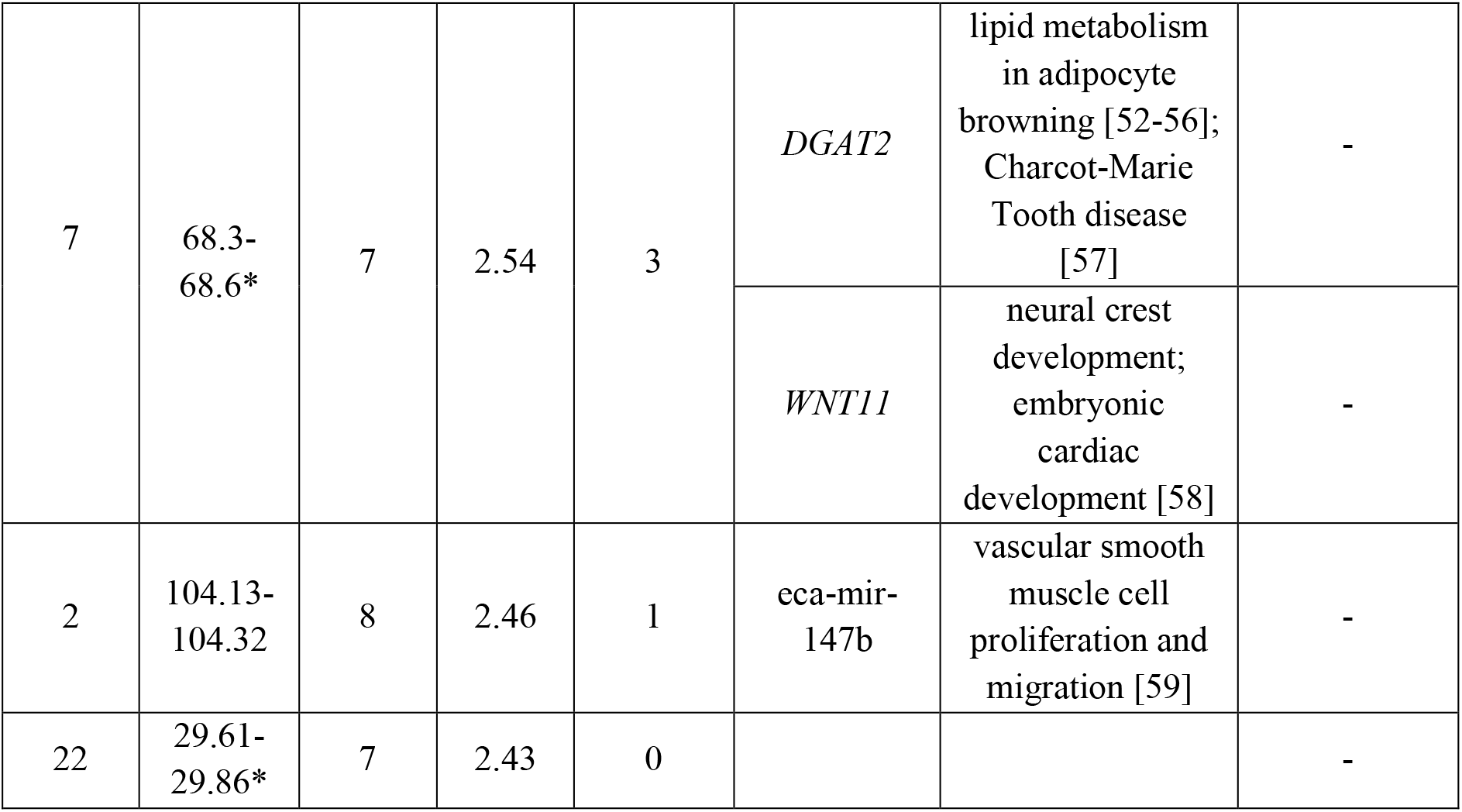
Selected genomic regions in Thoroughbreds containing ≥ 5 SNPs among the top 1% (480) SNPs ranked by the composite selection signal (CSS) value. Chr = ECA; Region = EqCab2.0. *Regions previously identified as being under selection in Thoroughbreds [26, 27].

| Chr | Region (Mb) | Top 1% SNPs (n) | Top CSS value | Cluster genes (n) | Candidate genes | Gene function | ROH overlap |
| --- | --- | --- | --- | --- | --- | --- | --- |
| 1 | 38.24-45.47* | 87 | 5.39 | 29 | *ZWINT* | neuropathic pain response [28-30] | ROH > 1Mb<br>ROH > 5Mb |
| 17 | 20.69-23.86* | 31 | 5.11 | 22 | *DLEU7* | growth traits [31]; overgrowth [32] | ROH > 1Mb |
|  |  |  |  |  | *KCNRG* | aortic root diameter [33] |  |
| 1 | 121.51-122.22* | 15 | 3.99 | 2 | *THSD4* | lung function [34] | - |
| 4 | 18.95-19.85 | 14 | 3.79 | 3 | *VWC2* | bone formation [35] | ROH > 1Mb<br>ROH > 5Mb |
| 30 | 24.6-25.26 | 18 | 3.49 | 2 | *ASPM* | cerebral cortex size [36] | - |
| 18 | 47.23-47.73 | 6 | 3.27 | 1 | *XIRP2* | cardiac hypertrophy [37] | ROH > 1Mb<br>ROH > 5Mb |
| 1 | 21.94-22.78 | 10 | 3.15 | 0 | *XPNPEP1* | response to stress [38]; behavioural hyperactivity [39] | - |
|  |  |  |  |  | *SORCS1* | insulin metabolism [40] | - |
| 1 | 178.8-179.86 | 16 | 3.0885<br>55 | 1 | *FKBP25* | protective response to ischemic injury [41] | ROH > 1Mb |
| 21 | 54.66-55.46 | 15 | 2.94 | 1 | *IRX1* | hip geometry [42]; bone mineral density [43] | ROH > 1Mb |
|  |  |  |  |  | *ADAMTS16* | high altitude adaptation [44] | - |
| 6 | 21.92-23.58 | 26 | 2.96 | 7 | *COL6A3* | collagen / tendon [45] [46] | ROH > 1Mb |
| 14 | 41.78-42.44* | 11 | 2.6016<br>78 | 1 | *FSTL4* | extinction of inhibitory avoidance memory; regulates BDNF [47-49] | - |
| 14 | 46.78-47.42* | 14 | 2.5416<br>32 | 3 | *MEGF10* | muscle cell proliferation; regulation of myogenesis [50, 51] | - |
| 7 | 68.3-68.6* | 7 | 2.54 | 3 | *DGAT2* | lipid metabolism in adipocyte browning [52-56]; Charcot-Marie Tooth disease [57] | - |
|  |  |  |  |  | *WNT11* | neural crest development; embryonic cardiac development [58] | - |
| 2 | 104.13-104.32 | 8 | 2.46 | 1 | eca-mir-147b | vascular smooth muscle cell proliferation and migration [59] | - |
| 22 | 29.61-29.86* | 7 | 2.43 | 0 |  |  | - |

We interrogated the CSS regions for candidate genes that may contribute to the Thoroughbred phenotype and identified genes with functions in behaviour, musculoskeletal, cardiac and respiratory function, conformation and metabolism (Table 1). Given that positive selection at a specific genomic locus tends to reduce (‘sweep’) variation across a larger region, it can be difficult to identify the gene under selection. Here, we identified plausible candidate genes driving selection based on the location of the highest-ranking SNPs in the selected region and by reviewing known biological functions of genes that we hypothesise may be selected for the Thoroughbred phenotype. Seven of the top 10 ranked SNPs were located in the top ranked region on ECA1 and the top three SNPs spanned 95 kb closest to the ZW10 interacting kinetochore protein gene (*ZWINT*). ZWINT (also known as SIP30) is abundantly expressed in the brain, modulates neurotransmitter release and functions in the mediation of peripheral nerve injury-induced neuropathic pain [28–30]. In rodents ZWINT influences pain-evoked emotional responses [60] and since human athletes have been reported to have a greater ability to tolerate pain than normally active controls [61], it is intriguing to speculate that ZWINT may be involved in the equine response to exercise-induced pain. There were particularly long ROH in this region; 31 of the 305 stallions had ROH >16 Mb between ECA1: 37.8-76.3 Mb (S1 Table); the longest ROH spanned almost 40Mb.

We identified other neurological/behaviour associated genes in the selected regions including the follistatin like protein 4 gene (*FSTL4*), the abnormal spindle microtubule assembly gene (*ASPM*) and the X-prolyl aminopeptidase 1 gene (*XPNPEP1*). Knockdown of *FSTL4* in mice results in the extinction of inhibitory avoidance memory indicating its involvement in synaptic plasticity and memory formation. Interestingly it functions by directly interacting with the exercise-induced brain derived neurotrophic factor (BDNF) [62, 63] which decreases in response to chronic stress [64]. *ASPM* is a major determinant of the size of the cerebral cortex in mammals [36, 65] that plays a key role in memory, attention, perception and awareness. Knockdown of *XPNPEP1*, which encodes an important downstream regulator of the stress response [38], in mice results in enhanced locomotor activity and impaired contextual fear memory [39]. An emerging theme in equine transcriptomics and genomics research suggests a link between the exercise phenotype and behavioural plasticity. For example, in the skeletal muscle transcriptome response to exercise training, neurological processes were the most significantly over-represented gene ontology (GO) terms, with the top three ranked GO terms being *Neurological system process* (*P* = 4.85 × 10^−27^), *Cognition* (*P* = 1.92 × 10^−22^) and *Sensory perception* (*P* = 4.21 × 10^−21^) [66]. Furthermore, in genome-wide association (GWA) studies genes involved in behavioural plasticity are the most strongly associated with economically important traits in racing Thoroughbreds: precocity (early adaptation to racing) [67] and the likelihood of racing [19]. For Thoroughbred horses, behavioural plasticity enables adaptation to the rigours of an intense exercise training programme in an unnatural environment, with considerable variation in the abilities of horses raised in the same environment to adapt to stress. The presence of these genes in genomic regions under selection in the Thoroughbred population supports human-mediated adaptation of the Thoroughbred towards heightened awareness and the ability to learn and adapt to stress.

Genes for aesthetic physical phenotypes including height, stature, coat and plumage colour are commonly encountered in genomic regions under selection in domestic animal populations since they are easily identifiable [68–71]. Conformation characteristics are among the most discernible traits in horses and are the principal observable traits on which Thoroughbreds are selected. Here, the selection signal on ECA21 centred on the iroquois homeobox 1 gene (*IRX1*) and also the ADAM metallopeptidase with thrombospondin type 1 motif 16 gene (*ADAMTS16*), which has been shown to be associated with hip geometry in humans [43]. While *ADAMTS16* has been identified among selection signals for high altitude adaptation in pigs [44], *IRX1* is associated with bone mineral density in humans [43] and influences chondrocyte differentiation and may therefore contribute to joint flexibility [72]. In horses, genes with functions in bone mineral density have been associated with measurements of joint angles [42]. Since the angle of the pelvis is a major determinant of physical conformation in Thoroughbreds and is associated with injury and performance [73], we hypothesise that the selection signature on ECA21 reflects the evolution of the Thoroughbred conformation phenotype.

Other genes relating to musculoskeletal form and function were identified among selected regions. For example, the deleted in lymphocytic leukemia 7 gene (*DLEU7*) is associated with growth traits in chickens [31] and overgrowth in humans [32] and may therefore contribute to stature in Thoroughbreds. In bone physiology, the von Willebrand factor C domain containing 2 gene (*VWC2*), a bone morphogenic protein, promotes bone formation, regeneration and healing [35, 74]. In the context of Thoroughbred musculature, the multiple EGF like domains 10 gene (*MEGF10*) controls muscle cell proliferation and is involved in the regulation of myogenesis [50, 51] and the collagen type VI alpha 3 chain gene (*COL6A3*) plays a major role in the maintenance of strength of muscle and connective tissue [45]. *COL6A3* is one of three genes encoding components of collagen VI, which in the horse is expressed in developing cartilage [46]. Also, collagen VI disruption in horses has been associated with osteochondrosis, a common developmental orthopaedic disease with a major economic impact in the Thoroughbred industry [75]. *COL6A3* may also be relevant to the muscle metabolism phenotype since its expression in adipocytes is associated with insulin resistance and obesity [76, 77]. Mutations in the diacylglycerol O-acyltransferase 2 gene (*DGAT2*) mutations cause Charcot-Marie Tooth disease in humans [57] and the DGAT2 protein also functions in lipid metabolism [52–56].

### Polygenic adaptation in the Thoroughbred

There are likely numerous endophenotypes on which selection acts to generate the athletic phenotype. Adaptation driven by selective sweeps at a number of key genomic loci is likely to be important; however, modest allele frequency changes at multiple loci are also expected to occur as a consequence of polygenic adaptation [78, 79]. Therefore, in addition to ‘core’ genes that are critical to the phenotypic outcome, highly granular additive genetic variation—essentially encompassing the entire genome—combined with epistatic interactions differing across cell types, may largely shape the phenotype [80]. Considering this, we performed an enrichment analysis to identify functional processes over-represented among the set of 387 of the 462 genes contained within the putative selection regions that mapped to the Ingenuity Pathway Analysis (IPA) database. The top canonical pathway was *Airway inflammation in asthma* (S5 Table). Manual curation of the top 50 canonical pathways points to a process of polygenic adaptation among functional modules in cardiovascular signalling, cellular growth, proliferation and development, cellular immune response, cellular stress and injury, metabolic pathways (fatty acid and lipid degradation/biosynthesis), neurotransmitters and other nervous system signalling and organismal growth and development (S6 Table). Our results support the development of the Thoroughbred phenotype via a contribution from major allele frequency shifts at ‘core’ genes contributing to behavioural, metabolic and conformation traits and genome-wide changes in functional modules that shape a range of exercise-relevant physiological adaptations.

### Concluding remarks

We report here a highly significant increase in inbreeding in the global Thoroughbred population during the last five decades, which is unlikely to be halted due to current breeding practices. Inbreeding results in mutational load in populations that may negatively impact on population viability. ‘Genetic rescue’ of highly inbred populations may be possible by the introduction of genetically diverse individuals [81]; however, rescuing genetic diversity in the Thoroughbred will be challenging due to the limitations of a closed stud book. Furthermore, the population has a small effective population size (*N*_e_) and a limited numbers of stallions have had a disproportionate influence on the genetic composition of the Thoroughbred; 97% of pedigrees of the horses included here feature the ancestral sire, *Northern Dancer* (b. 1961) and 35% and 55% of pedigrees in EUR and ANZ contain *Galileo* (b. 1998) and *Danehill* (b. 1986), respectively (S1 Text).

Pedigree data can be useful to illustrate broad trends in breeding practices, but since pedigree-based estimates of inbreeding and relatedness have poor correlations with estimates using genomic methods [27, 82–85] relying on pedigree alone for outcrossing is likely to be inefficient in reversing the trends observed here (S1 Text, S19 Figure). Directives to prevent over-production from popular sire lines and the global movement of stallions that are distinct from the local population of mares may act to maintain and increase genetic diversity in the population. However, given the limited diversity in current Thoroughbred pedigrees, genomics-based measures using high-density genome-wide SNP information and a large reference population are likely to offer the best opportunity to slow and reverse the potential effects of inbreeding. The introduction of an industry-mediated longitudinal programme of genomics-based monitoring of inbreeding and the implementation of guidelines and strategies for genome-enabled breeding that are comparable to methods used in other domestic species, will contribute to promoting economic gain and safeguarding the future of the breed.

## Methods

### Ethics statement

Animals used in this study were submitted for genetic analyses and consent for use of samples in research was obtained during the sample submission process.

### Sampling and population assignment

The following data was collected for *n* = 10,118 Thoroughbred (TB) horses: horse name, sire, dam, sex, year of birth and country of birth (S7 Table). Based on country of birth, horses were assigned to the following geographic regions: Europe/Middle East (EUR), Australasia (ANZ), North America (NAM), South Africa (SA) and Japan (JAP). An additional set of *n* = 84 horses from four breeds that were chosen to represent putative TB founder populations included *n* = 20 Akhal Teke (AKTK), *n* = 24 Arabian (ARR), *n* = 23 Moroccan Barb (MOR) and *n* = 17 Connemara pony (CMP). The ARR and AKTK sample data used were obtained from publicly available equine Illumina SNP50 Beadchip genotype data [3].

### Assembly of comparative SNP data set, quality control and filtering of SNPs

DNA was isolated from blood or hair samples and genotyped using the Illumina EquineSNP50 BeadChip (SNP50), the Illumina EquineSNP70 BeadChip (SNP70) or the Affymetrix Axiom™ Equine 670K SNP genotyping array (SNP670). Only animals and SNPs with a genotyping rate > 95% were included with a minor allele frequency (MAF) threshold > 0.05 applied. A set of 48,896 autosomal SNPs derived originally derived from the SNP50 and SNP70 arrays was used for the analysis. This SNP set was extracted from the genotype data from each of the three platforms.. SNPs that failed quality control in < 5% of samples or were not present on one of the array platforms were imputed using the software program BEAGLE (version 3.3.2; [86, 87]). For 10 horses genotyped using both the SNP50 and SNP70 arrays and 10 different horses separately genotyped using the SNP70 and SNP670 array post-imputation concordance was greater than 99%.The TB dataset (*n* =10,118) comprising of the set of 48,896 SNPs was pruned using the PLINK software suite V1.9 (http://pngu.mgh.harvard.edu/purcell/plink/; [88]) -indep function with the following parameters: a five-step sliding window size of 50 with a VIF threshold of 50 where the VIF is 1/(1-R^2). This pruned set of 9,212 SNPs was used for principal component analysis (PCA) within the Thoroughbred population and for the calculation of individual inbreeding coefficients (*F*) as outlined below.

The Thoroughbred population was compared to founder populations using PCA and composite selection signature (CSS) analysis. We randomly selected *n* = 229 elite horses (CPI > 5, *i.e.* earned more than five times the average; CPI is a class performance index defined by the American Jockey Club) from the TB dataset (*n* = 10,118). Genomic relationships among horses were estimated using autosomal identity by descent (pi-hat) values in PLINK v1.9 [89]. After removing individual horses with pi-hat > 0.25, 150 horses (*n* = 23 MOR, *n* = 17 CMP, and *n* = 110 TBE) remained. Then, the newly genotyped data was merged with publicly available data for AKTK (*n* = 20) and ARR (*n* = 24) horses which was pre-processed to include only individuals and SNPs with a genotyping rate > 95% and with MAF > After a final round of quality control (MAF > 0.01 and genotyping rate > 0.95) on the combined data set, 194 horses with 31,722 SNPs remained for the mixed breed PCA and CSS analyses.

### Stallion genotype reconstruction

To increase the representation of prominent stallions, genotypes (~ 46,000 autosomal SNPs) were reconstructed for horses where genotypes were available for *n* ≥20 progeny of an individual stallion (S8 Table). An adaptation of the method described by Gomez-Raya [90] was used to infer genotypes. The main difference here was that population genotype frequencies were included allowing accurate reconstruction with just 20 offspring. If all offspring do not share one allele at a locus the sire must be heterozygous at the locus. If all offspring do share one allele at a locus the sire may be either heterozygous at the locus or homozygous for the shared allele. However, the probability of the sire being homozygous or heterozygous can be calculated based on the proportion of the different genotypes in the offspring and the observed proportion of each genotype in the population. The chi-square statistic was calculated using observed and expected offspring allele frequencies for each of the three possible sire genotypes. In order to compare the three sire genotype possibilities and assign a relative probability to each, the chi-square test statistics were first converted to likelihood scores. To do this the density of a chi-square distribution was taken at each of the three points (the density at each point represents the likelihood of the sire having that genotype, given the observed offspring genotypes). The density values were then divided by the sum of the three densities to normalize. Normalizing these three likelihoods by their sum gave the relative likelihood of each sire genotype and the genotype with the maximum likelihood was assigned as the sire genotype.

### Principal component analysis (PCA)

PCA was conducted using smartPCA from the EIGENSOFT package (version 4.2) [91]. To include all the horses in the analysis, an option, outliersigmathresh, was set as 10. All other parameters were set to default values. Principal components (PCs) were plotted with data points colour-coded based on each horse’s geographic region of origin. A series of four separate analyses were performed to investigate Thoroughbred population sub-structure. For mixed breed analysis (1) the set of 31,722 SNPs were used while the pruned SNP set (9,212 SNPs) was used for the within Thoroughbred analyses (2-4).

1. Thoroughbreds and putative founder breeds comprised a set of *n* = 110 randomly selected elite TB and *n* = 20 AKTK, *n* = 24 ARR, *n* = 23 MOR and *n* = 17 CMP.
2. Global Thoroughbred population comprised a set of all Thoroughbred samples (*n* = 10,118) in the data set.
3. Global sire lines comprised a set of *n* = 305 globally distributed prominent stallions.
4. TB geographic regions – EUR (*n* = 4,099), ANZ (*n* = 2,733) and NAM (*n* = 1,992).

### Pedigree analysis

Ten generation pedigree data was ascertained for *n* = 7,262 Thoroughbred horses (www.pedigreequery.com)

### Inbreeding estimates

Individual inbreeding coefficients (*F*_IS_) for each horse were calculated in PLINK based upon reduction in heterozygosity relative to Hardy-Weinberg expectation (–het Genomic inbreeding was also evaluated by identifying runs of homozygosity (ROH) with the --homozyg command. ROH were defined as tracts of homozygous genotypes that were >1 Mb in length identified for one SNP per 1000 kb on average and two consecutive SNPs < 1000 kb apart. No more than two missing genotypes and one heterozygous genotype were allowed. The following parameters were set: -- homozyg; -- homozyg-kb 1000; --homozyg-snp 30; -- homozyg-gap 1000; -- homozyg-window-het 1; --homozyg-window-snp 30; -- homozyg-density 1000; -- homozyg-window-missing 2 to identify runs of homozygosity spanning at least 1 Mb. The analysis was also run with the parameter homozyg-kb increased to 5000 to identify runs of homozygosity of 5 Mb or greater. First, the individual sum of total ROH per animal was calculated. The *F*_ROH_ statistic proposed by McQuillan *et al.* [92] was then calculated, whereby the total length of ROH covering an individual animal’s genome (L_ROH_) is divided by the length of the autosomal genome (L_AUTO_); *F*_ROH_ = L_ROH_/L_AUTO_. Here, we used the length of the equine autosomal genome as 2,242,960 kb (www.ncbi.nlm.nih.gov/genome/145?genome_assembly_id=22878).

The annual mean, SE, SD and CI were calculated based on the horse’s year of birth for both measures of inbreeding (S16-S17 Table). Following a preliminary analysis, the pre-1996 samples were excluded from further analysis as sample distribution pre-1996 was low. The post-1995 data is of the most interest as this coincides with the introduction of “big books” for stallions; i.e. large numbers of mares bred. The shuttling of stallions to Australia from Europe for dual hemisphere breeding seasons peaked in 2001. A summary of the year of birth, region of birth and sex of the horses used in these analyses is provided in S7 Table. A pair-wise Student’s t-Test was used to compare measures of inbreeding across the main geographic regions represented in the data set; Europe/Middle East (EUR), Australasia (ANZ) and North America (NAM).

A linear model was used to assess the relationship between year of birth and inbreeding for the global population and for each of the geographic regions. Mean annual inbreeding values were used in the regression model (S9 Table, S10 Table). Pearson correlation was run to assess the relationship between inbreeding and racing performance defined by handicap (Timeform) rating. Significance was calculated by testing the null hypothesis of no linear correlation (r = 0).

### Effective population size (*N*_e_) estimates

The pruned set of 9,212 SNPs was used for the calculation of effective population size (*N*_e_). To estimate *N*_e_, plink-formatted data was first converted to GENEPOP format using PGDSpider 2.1.1.5 [93]. Then the LD method in NeEstimator2x [94] was used to calculate *N*_e_ using the converted GENEPOP data as input. *N*_e_ was calculated for global thoroughbreds and individual regions (ANZ, EUR,NAM and SAF). Year of birth for the horses used for *N*_e_ calculation were restricted to 2013 -2017.

### Composite selection signals (CSS)

The composite selection signals (CSS) method was used to identify signatures of selection in elite Thoroughbred horses (*n* = 110) using a mixed set of non-Thoroughbred horses representing putative founder populations (MOR, *n* = 23; CMP, *n* = 17; ARR, *n* = 24; AKTK, *n* = 20) as the comparator population.

The CSS approach was developed to investigate genomic signatures of selection and has been successful at localizing genes for monogenic and polygenic traits under selection in livestock [24, 25, 95]. The CSS uses fractional ranks of constituent tests and does not incorporate the statistics with *P* values, allowing a combination of the evidence of historical selection from different selection tests. For the present study, the CSS combined the fixation index (*F*_ST_), the change in selected allele frequency (Δ*SAF*) and the cross-population extended haplotype homozygosity (*XP-EHH*) tests into one composite statistic for each SNP. *F*_ST_ statistics were computed as the differentiation index between the population/s of interest (*i.e.* selected) and the contrasting/reference population/s (*i.e.* non-selected). *XP-EHH* and Δ*SAF* statistics were computed for the selected population(s) against the reference population. The CSS were computed as follows:

1. For each constituent method, test statistics were ranked (1, …, *n*) genome-wide on *n* SNPs.
2. Ranks were converted to fractional ranks (r′) (between 0 and 1) by 1/ (*n* + 1) through *n* / (*n* + 1).
3. Fractional ranks were converted to z-values as z = Φ-1(r′) where Φ-1(·) is the inverse normal cumulative distribution function (CDF).
4. Mean z scores were calculated by averaging z-values across all constituent tests at each SNP position and *P*-values were directly obtained from the distribution of means from a normal N (0, m^−1^) distribution where m is the number of constituent test statistics.
5. Logarithmic (−log_10_ of *P*-values) of the mean z-values were declared as CSS and were plotted against the genomic positions to identify the significant selection signals.
6. To reduce spurious signals, the individual test statistics were averaged (smoothed) over SNPs across chromosomes within 1 Mb sliding windows.

According to the approach proposed by Randhawa *et al*. [24], significant genomic regions were defined as those that harbour at least one significant SNP (top 0.1%) surrounded by at least five SNPs among the top 1%. Here, we relaxed the stringency to define significance as regions harbouring at least five SNPs among the top 1% since the numbers of regions would otherwise be small (i.e. ~48 SNPs). Also, since linkage disequilibrium extends up to 0.4 Mb in the Thoroughbred [96], we considered 1 Mb sliding windows reasonable in this population. Therefore, SNPs among the top 1% smoothed CSS values within the sliding windows were considered significant. Genes underlying the selection peaks (with at least one top 1%) as well as flanking regions (± 0.5 Mb) were identified by mapping to an annotated protein coding gene list downloaded from NCBI (accessed: 2018-05-17). These genes were then examined for evidence of functional significance.

### Gene Enrichment

Ingenuity^®^ Pathway Analysis (IPA^®^: Qiagen, Redwood City, CA, USA; release date June 2017) was used to perform an enrichment analysis to identify functional processes over-represented among the gene lists. Settings were such that the reference set was Ingenuity Knowledge Base (genes only); relationships to include were direct and indirect; interaction networks did not include endogenous chemicals. All other settings were default. *P* is reported as −log_10_ of the adjusted *P* value obtained with the Benjamini-Hochberg procedure [97]. Ratio denotes the ratio of genes specific to the pathway identified among selected regions in this study divided by the total number of the genes in this pathway designated by the Ingenuity Knowledge Base. Manual curation to group biological functions/pathways was performed using pathway information from https://targetexplorer.ingenuity.com.

## Supporting Information Text

**S1 Text:** Analysis of inbreeding and selection signatures in the Thoroughbred population

## Supporting Information Tables

**S1 Table** 31 stallions ROH at ECA1 region. Horses that were genotyped have been de-identified. Horses for which genotypes were imputed from progeny genotypes are identified.

**S2 Table:** Clusters identified with ≥5 SNPs among the top 1% based on the smoothed CSS statistic result (-log10P) for TB versus TB founder population comparison

**S3 Table:** The top 1000 SNPs ranked by the % individuals with SNP located within runs of homozygosity (ROH) of > 1mb

**S4 Table:** The top 1000 SNPs ranked by the % individuals with SNP located within runs of homozygosity (ROH) of > 5mb

**S5 Table:** Canonical pathways identified in the set of genes underlying selection peaks and flanking regions (± 0.5 Mb) for Thoroughbred (TB) versus TB founder population comparison using IPA analysis

**S6 Table:** 50 top canonical pathways identified in the set of genes underlying selection peaks and flanking regions (± 0.5 Mb) for Thoroughbred (TB) versus TB founder population comparison using IPA analysis by manual curation

**S7 Table:** Summary of distribution of horses based on region of birth and year of birth

**S8 Table:** List of imputed sires and number of progeny used for imputation, Dist is the genetic distance calculated by the pairwise identity by descent defined as (IBS2 +0.5*IBS1)/(n SNP pairs).

**S9 Table:** The annual mean, SE, SD and CI were calculated based on the horse’s year of birth for Inbreeding coefficient *F*.

**S10 Table:** The annual mean, SE, SD and CI were calculated based on the horse’s year of birth for *ROHs*

## Supporting Information Figures

**S1 Figure:** Thoroughbreds and breeds of origin PC1 v PC2

**S2 Figure:** Thoroughbreds and breeds of origin PC3 v PC4

**S3 Figure:** Global genetic variation in Thoroughbreds PC1 v PC2

**S4 Figure:** Global genetic variation in Thoroughbreds PC3 v PC4

**S5 Figure:** Stallions PC3 v PC4

**S6 Figure:** Within region genetic variation in Thoroughbreds – EUR PC1 v PC2

**S7 Figure:** Within region genetic variation in Thoroughbreds – EUR PC3 v PC4

**S8 Figure:** Within region genetic variation in Thoroughbreds – ANZ PC1 v PC2

**S9 Figure:** Within region genetic variation in Thoroughbreds – ANZ PC3 v PC4

**S10 Figure:** Within region genetic variation in Thoroughbreds – NAM PC1 v PC2

**S11 Figure:** Within region genetic variation in Thoroughbreds – NAM PC3 v PC4

**S12 Figure:** Global genetic variation of mean *F*_IS_ and *F*_ROH_ over time with standard error indicated by error bars.

**S13 Figure:** Regional genetic variation in annual mean *F*_IS_ and *F*_ROH_

**S14 Figure:** Linear regression fit for inbreeding by Year of birth in Australasia (ANZ)

**S15 Figure:** Linear regression fit for inbreeding by Year of birth in Europe (EUR)

**S16 Figure:** Linear regression fit for inbreeding by Year of birth in North America (NAM)

**S17 Figure:** Manhattan plot for CSS and smoothed CSS values, showing significant gene regions under selection for Thoroughbreds

**S18 Figure:** Manhattan plot of the distribution of runs of homozygosity (Greater than 1Mb in length and SNP located within ROH in > 20% of the population) in the Thoroughbred population.

**S19 Figure:** Comparison of pedigree-based estimates (x-axis) and the genomic estimate of relatedness (y-axis) of direct descendants of *Danehill*

