## Supplemental_Figures for "Genomic inbreeding trends in the global Thoroughbred horse population driven by influential sire lines and selection for exercise trait-related genes"

### S1 Figure
### Thoroughbreds and breeds of origin PC1vPC2

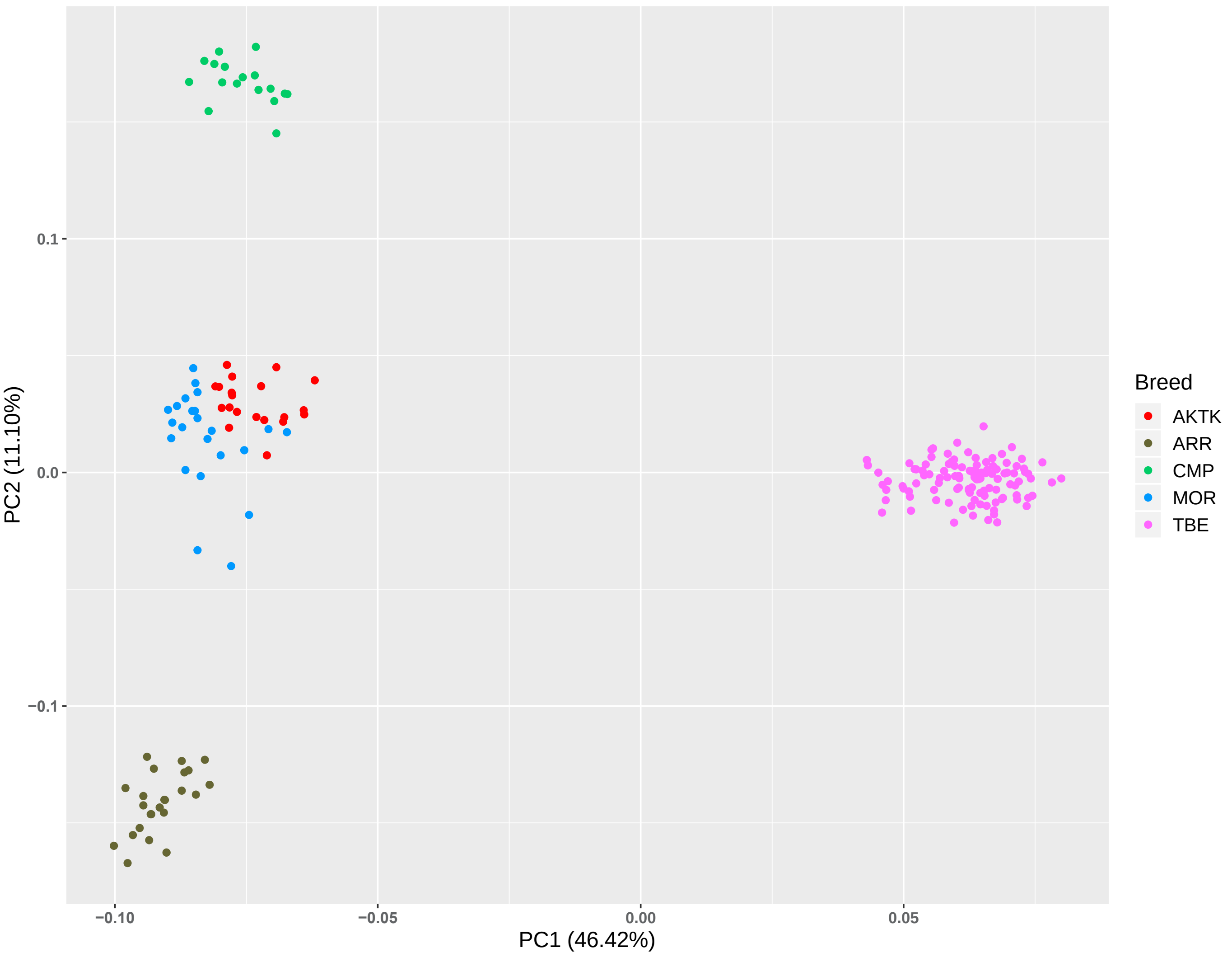

### S2 Figure
#### Thoroughbreds and breeds of origin PC3vPC4

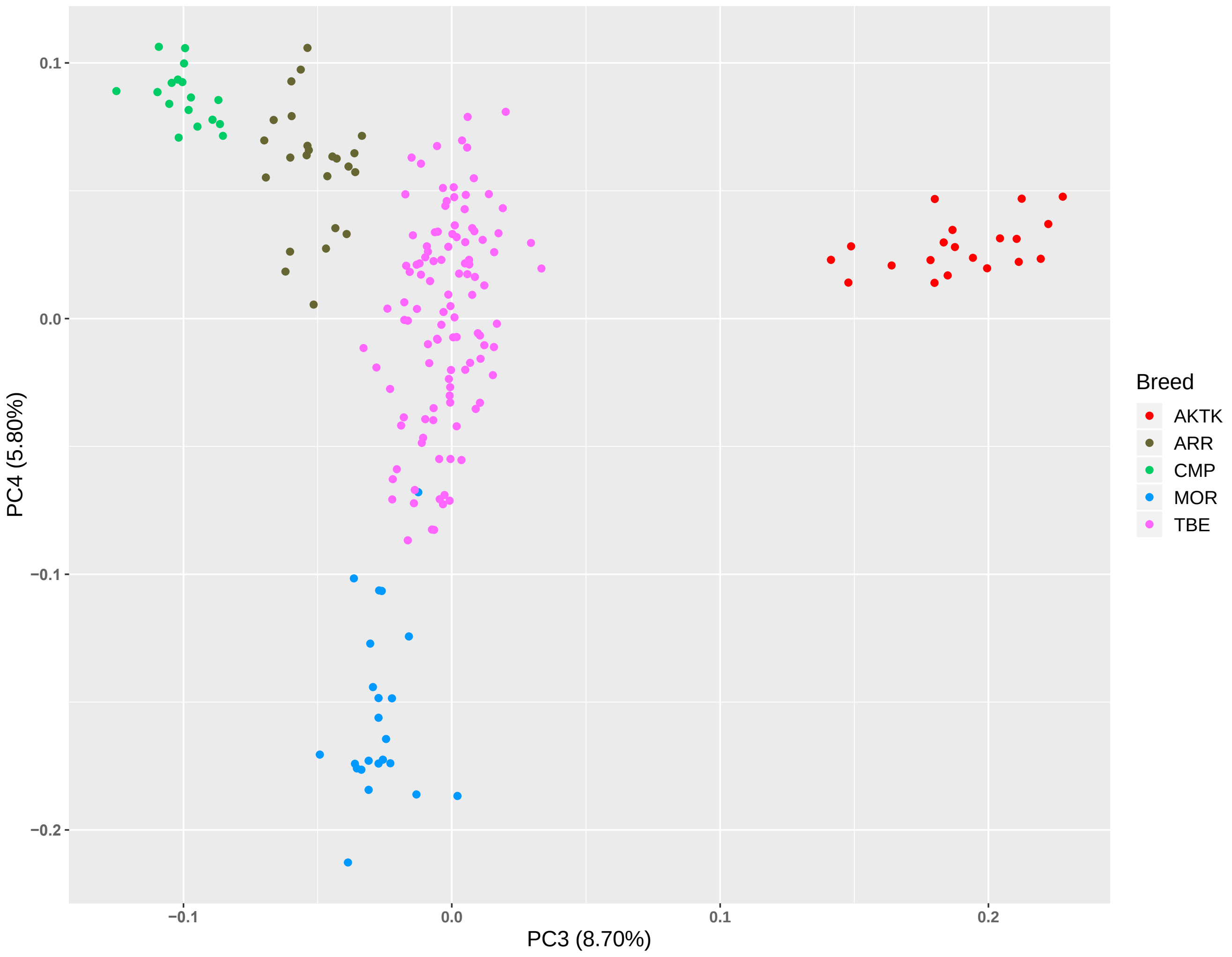

### S3 Figure
### Global variation in Thoroughbreds PC1 v PC2

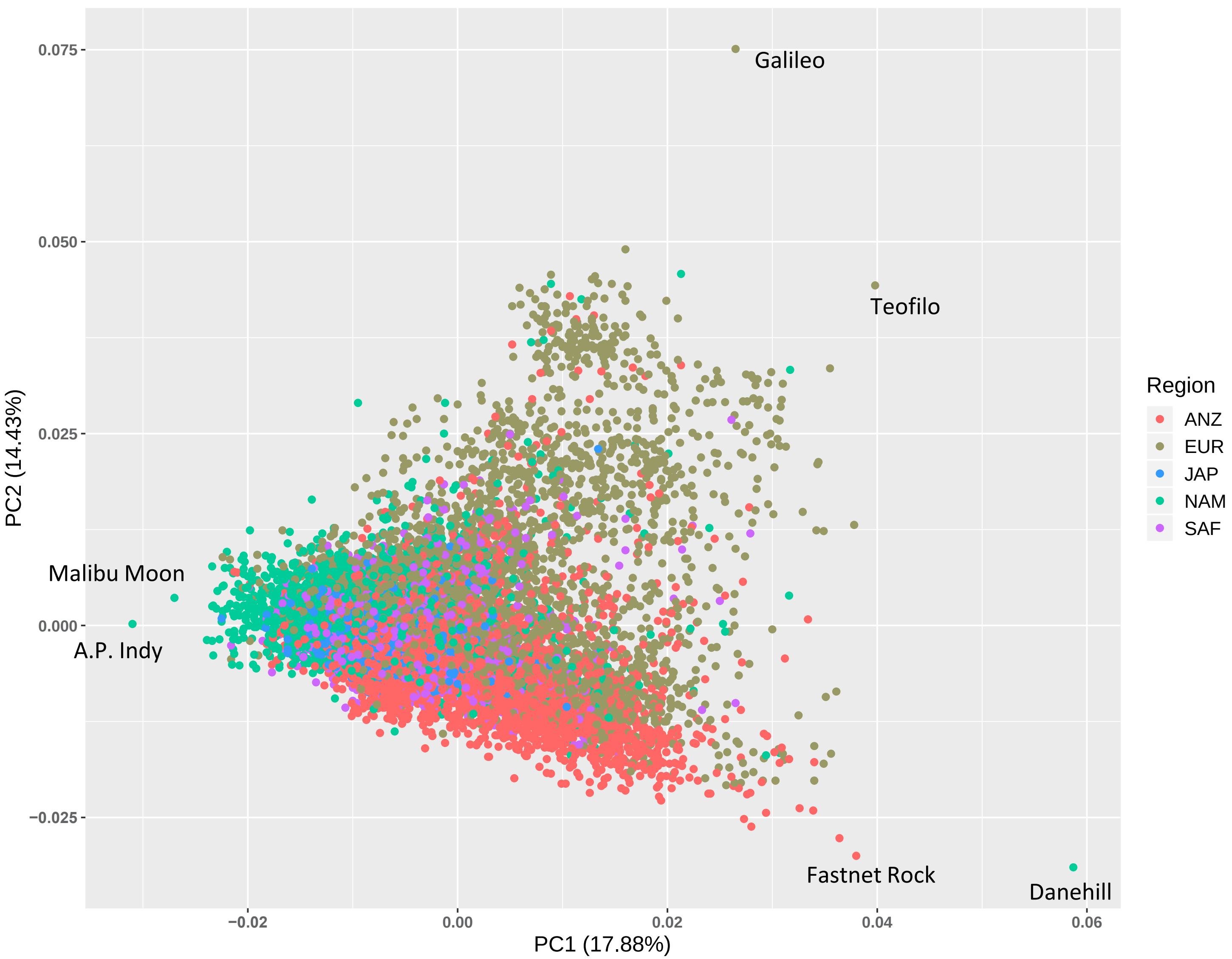

### S4 Figure
### Global variation in Thoroughbreds PC3 v PC4

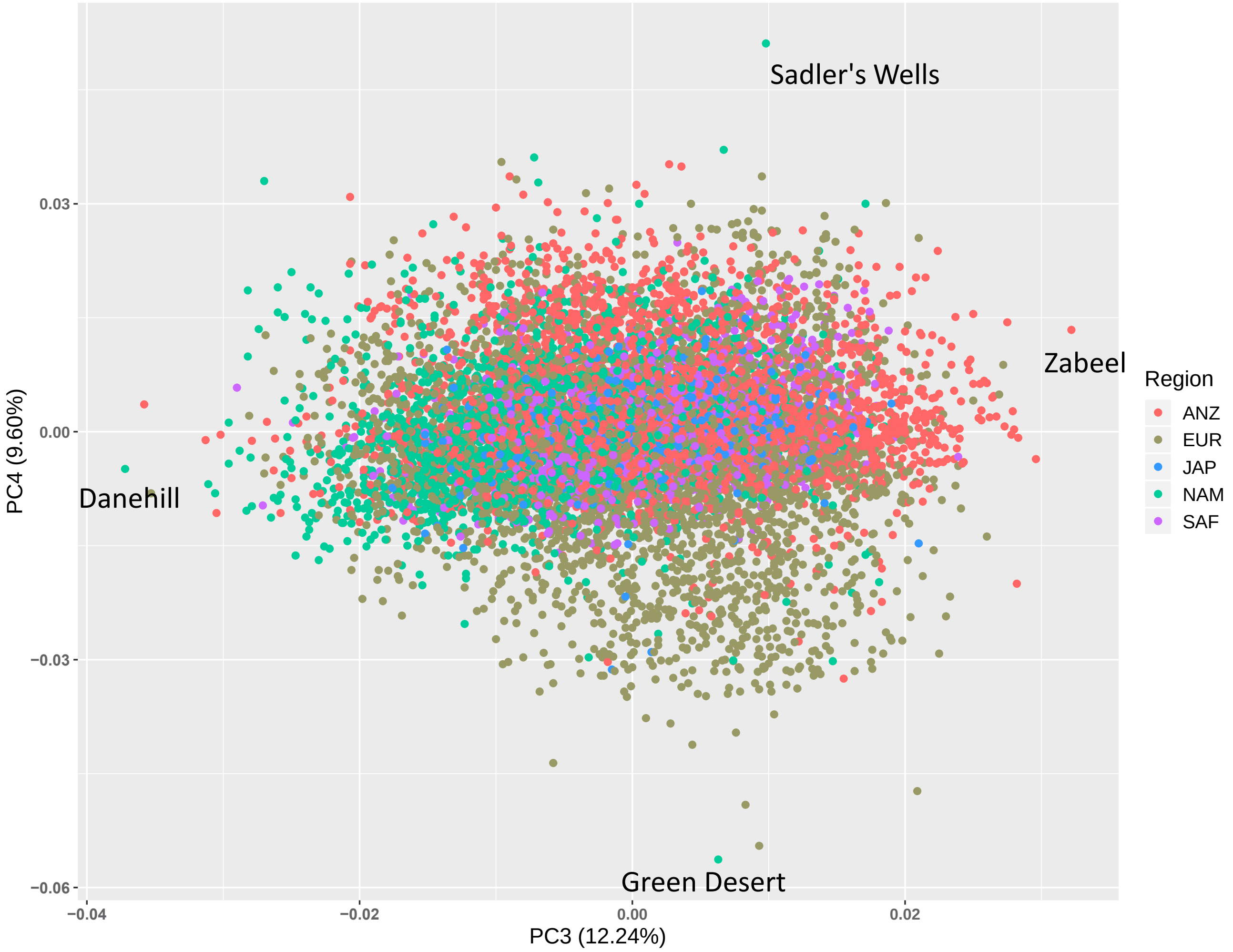

### S5 Figure
#### Stallions (n = 305) PC3vPC4

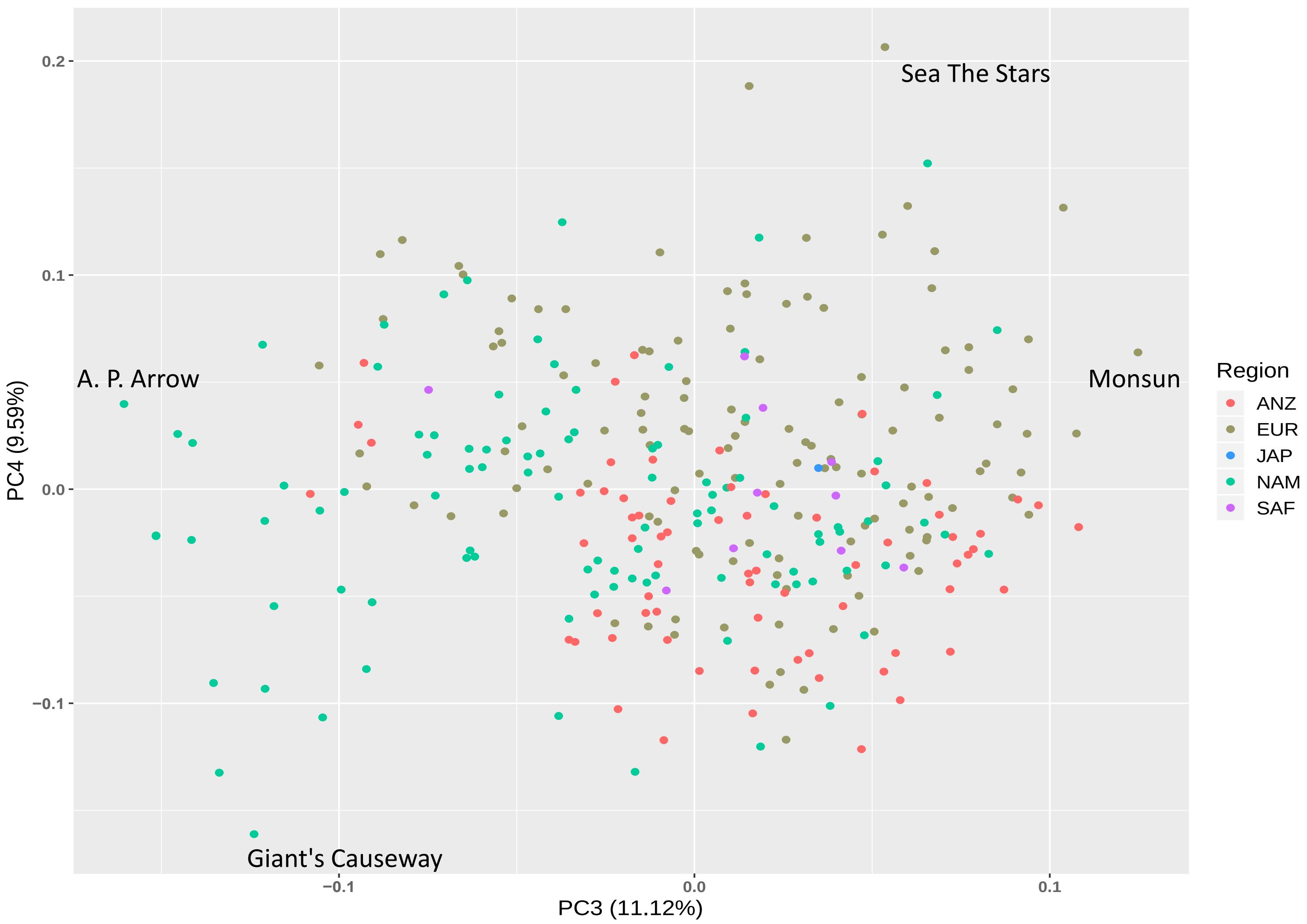

### S6 Figure
### Within region variation in Thoroughbreds – EUR
### PC1vPC2

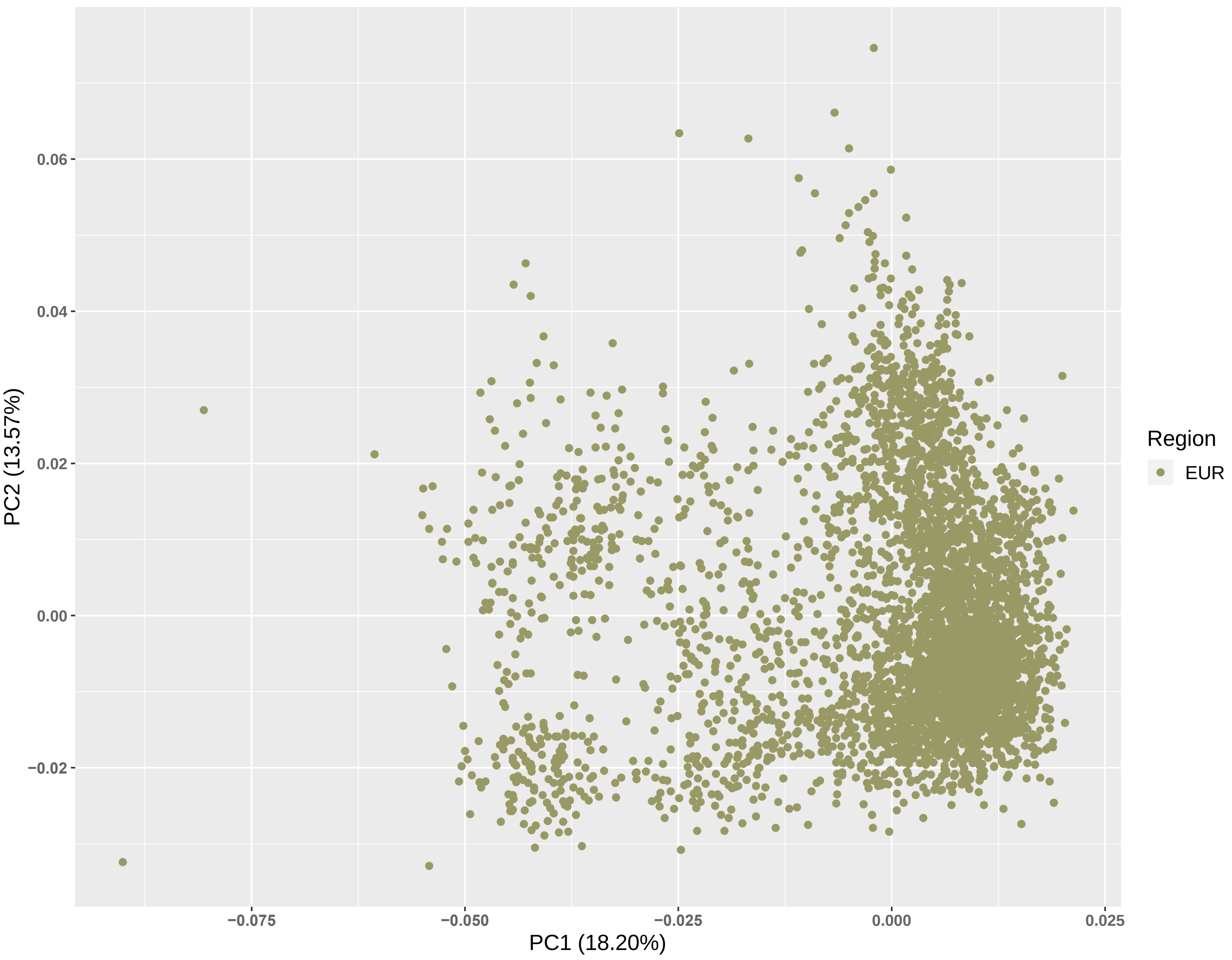

### S7 Figure
### Within region variation in Thoroughbreds – EUR
### PC3vPC4

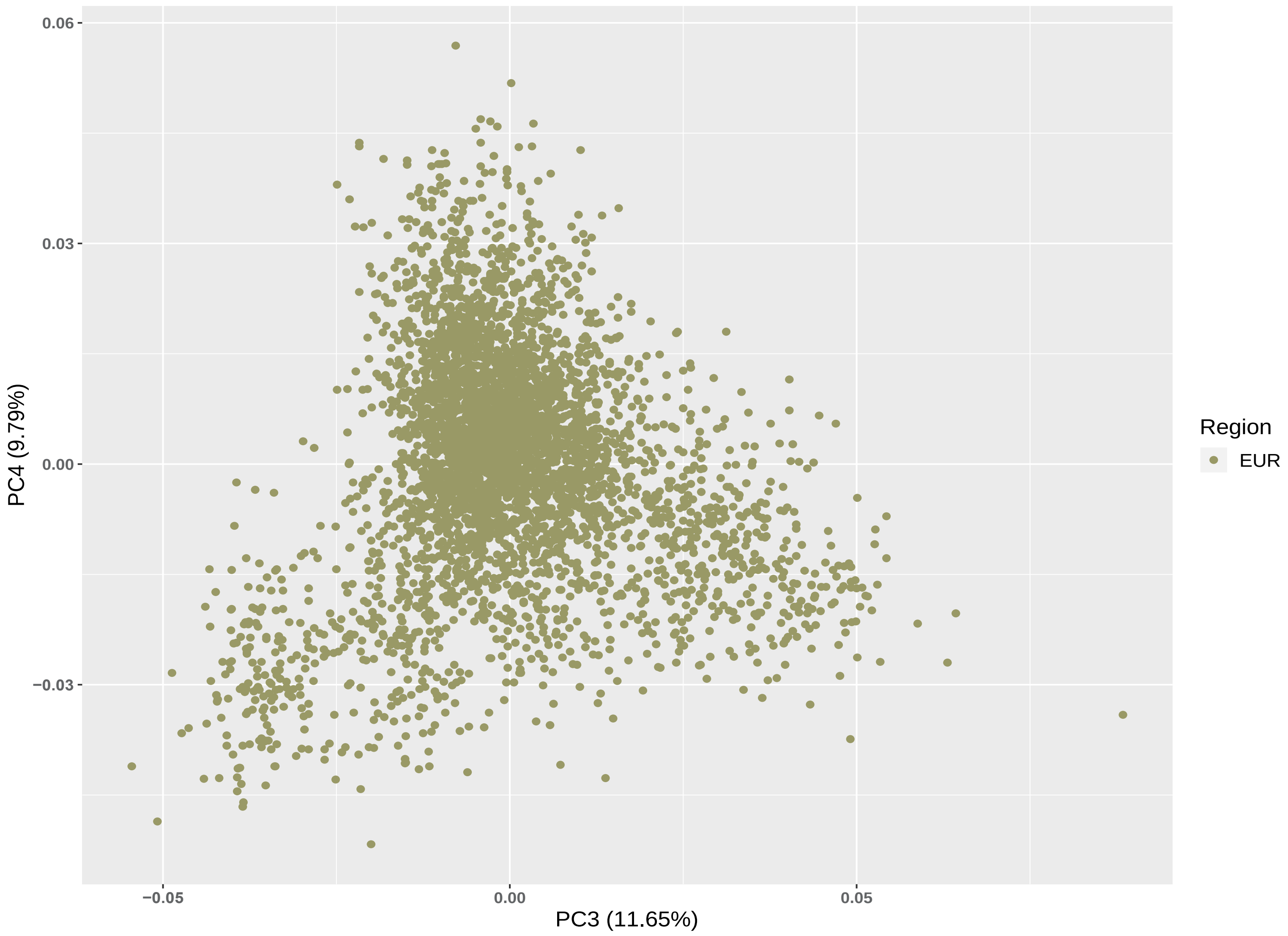

### S8 Figure
### Within region variation in Thoroughbreds – ANZ
### PC1vPC2

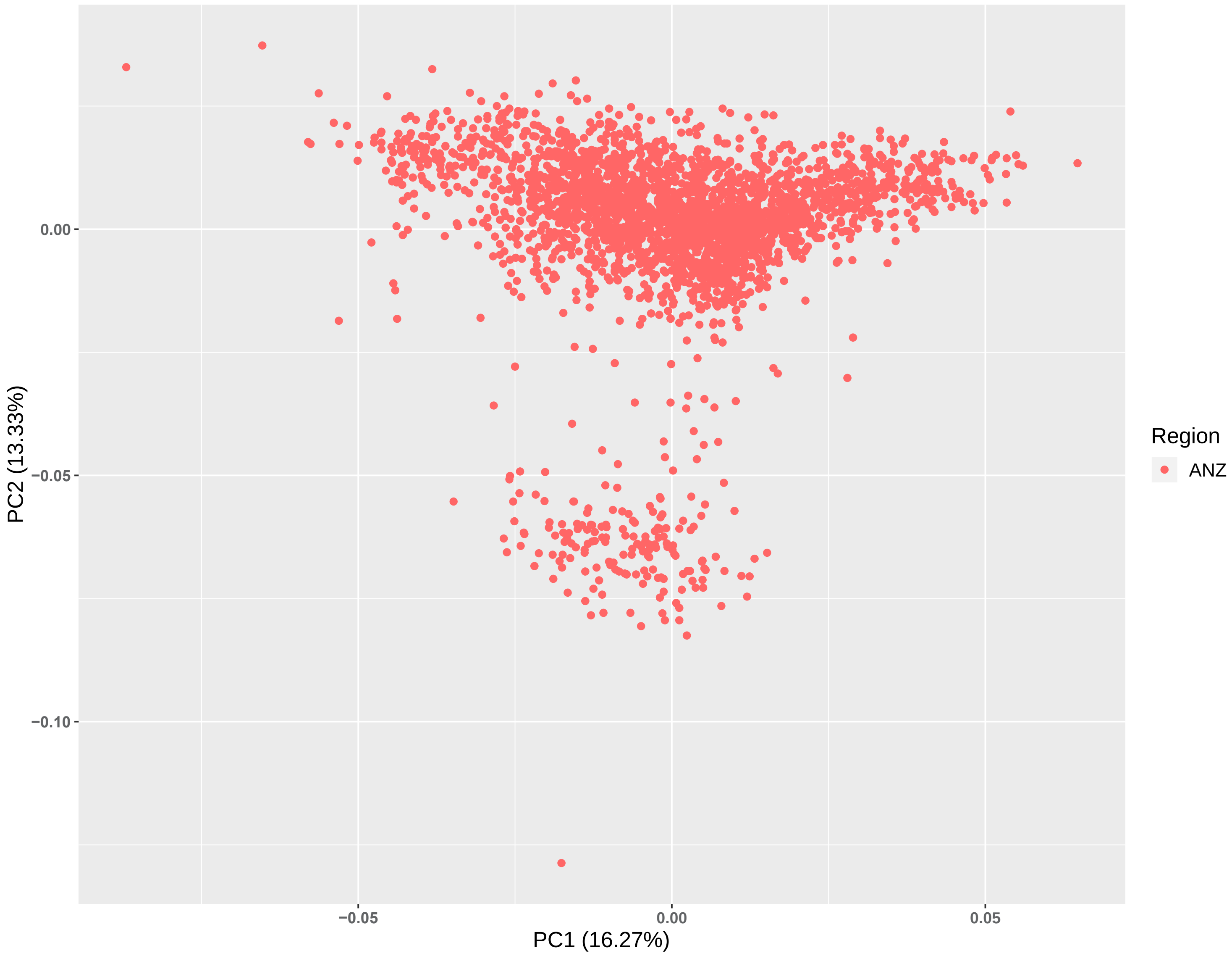

### S9 Figure
### Within region variation in Thoroughbreds – ANZ
### PC3vPC4

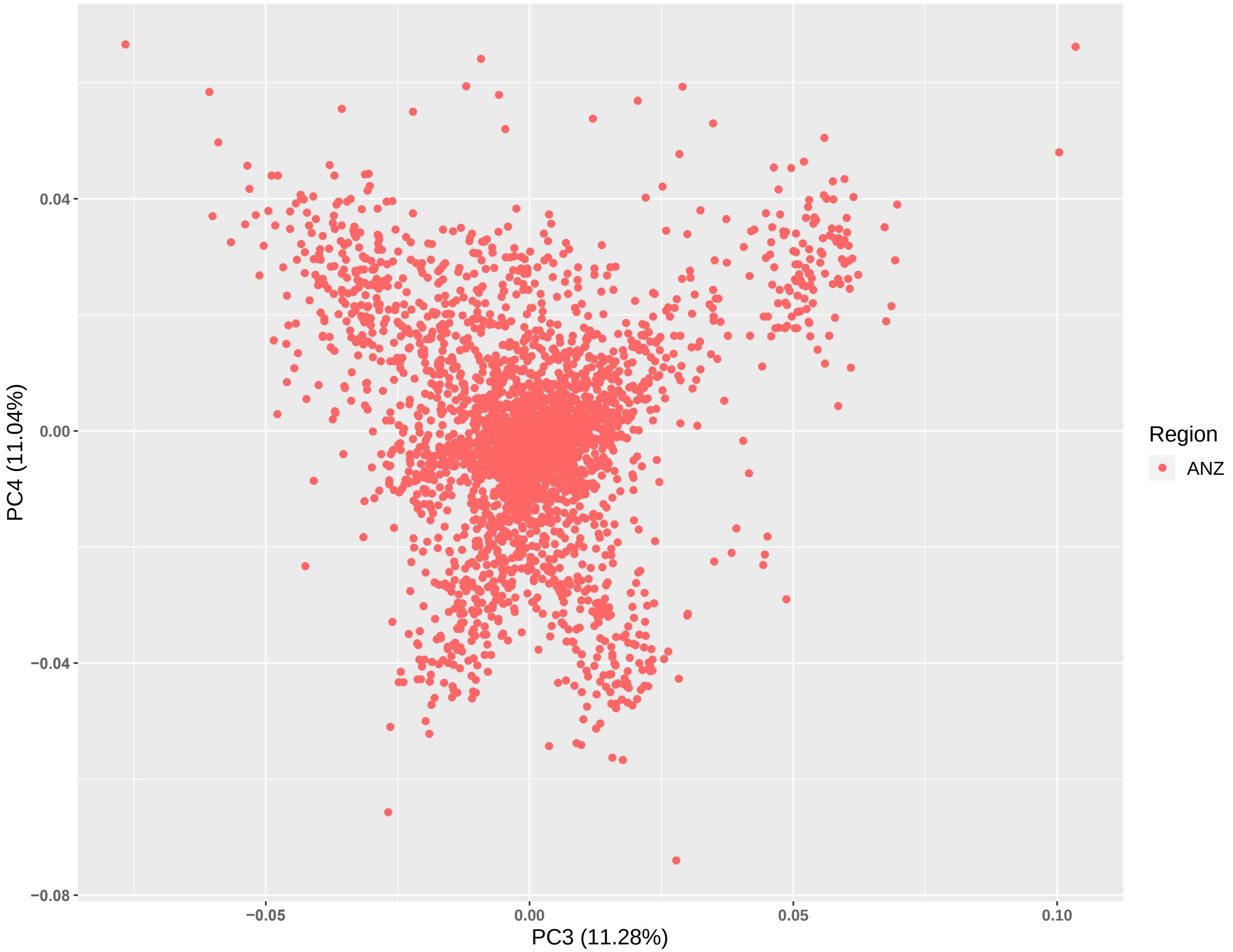

S10 Figure
Within region variation in Thoroughbreds – NAM
PC1vPC2

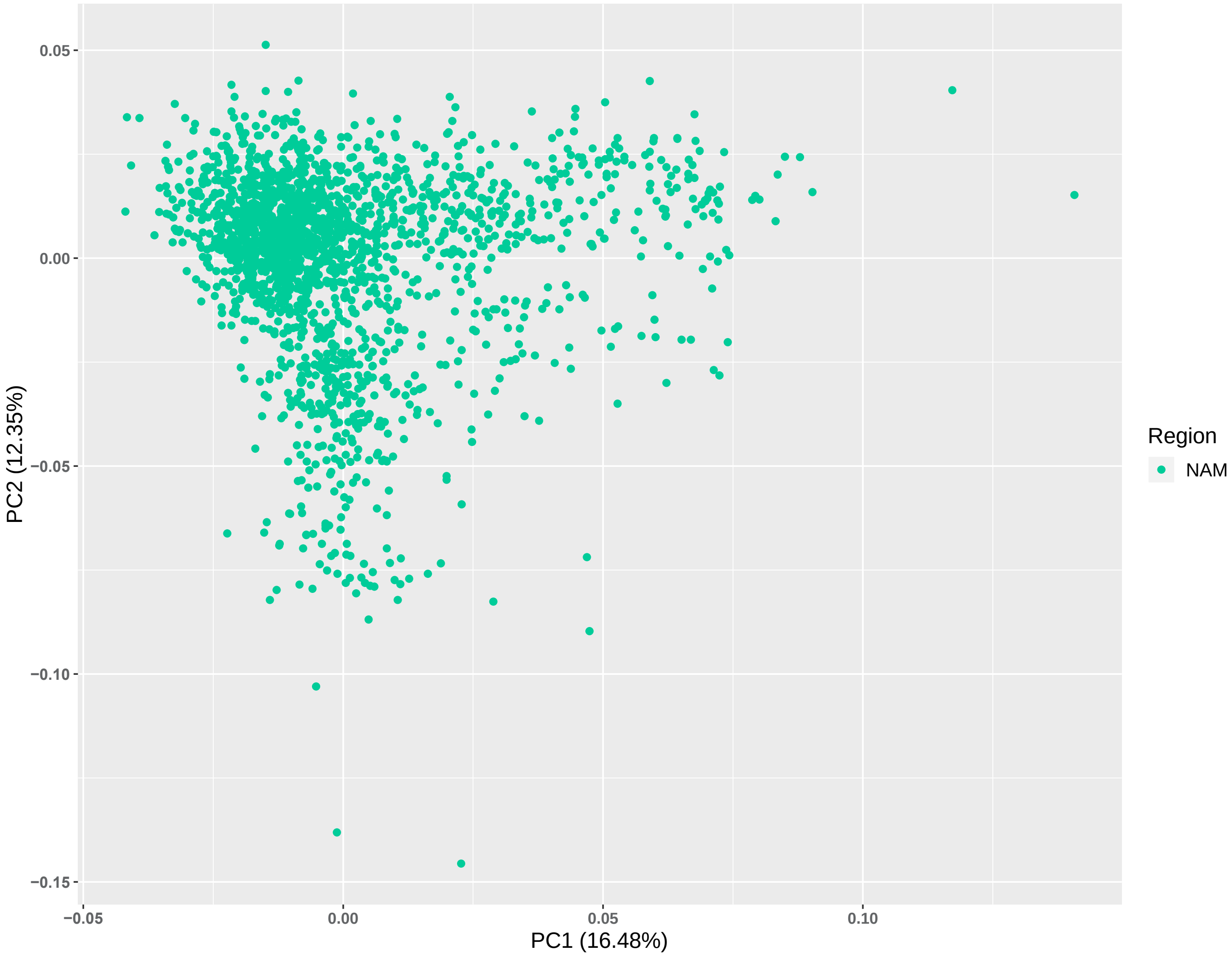

S11 Figure
Within region variation in Thoroughbreds – NAM
PC3vPC4

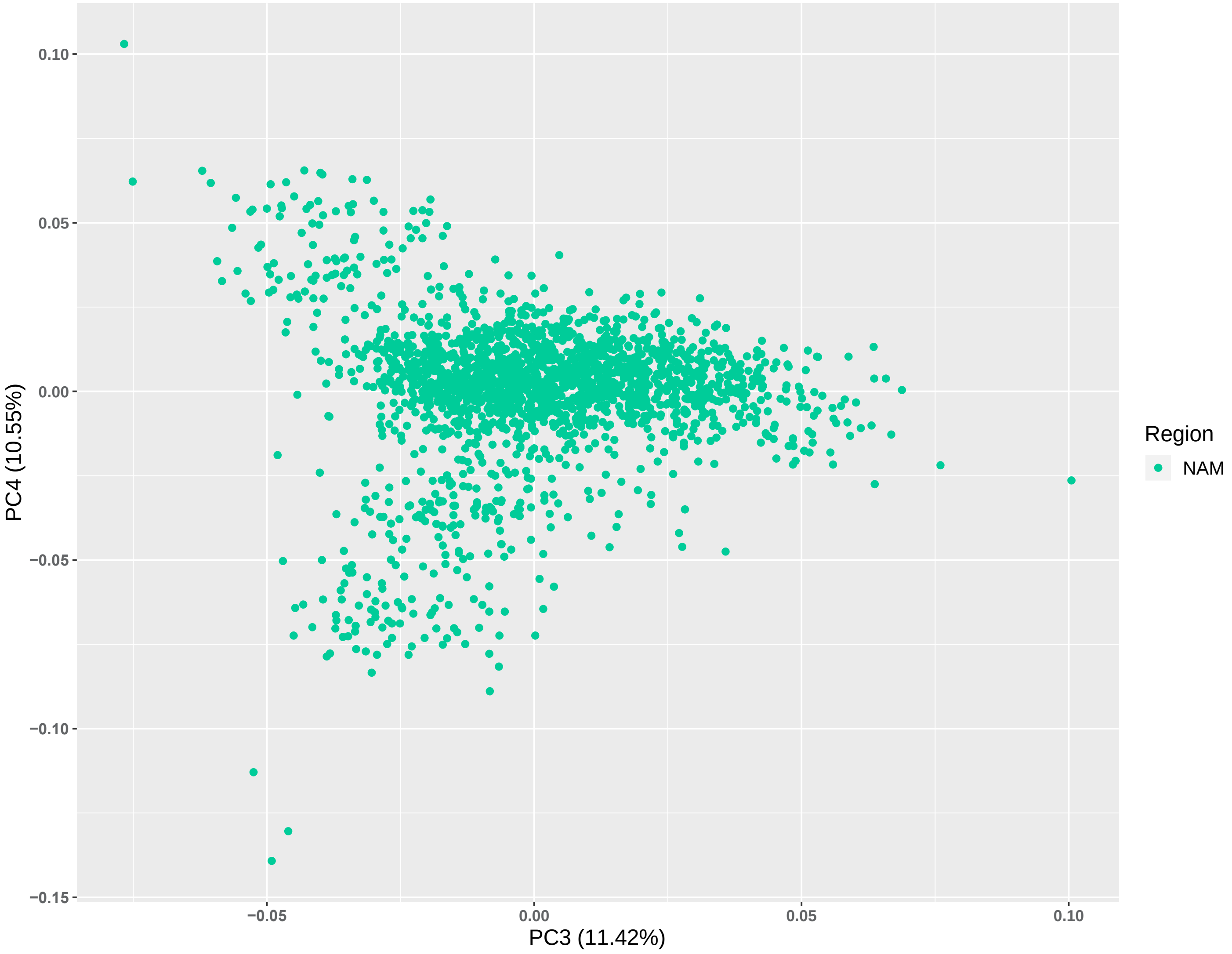

### S12 Figure
### Global variation in inbreeding over time

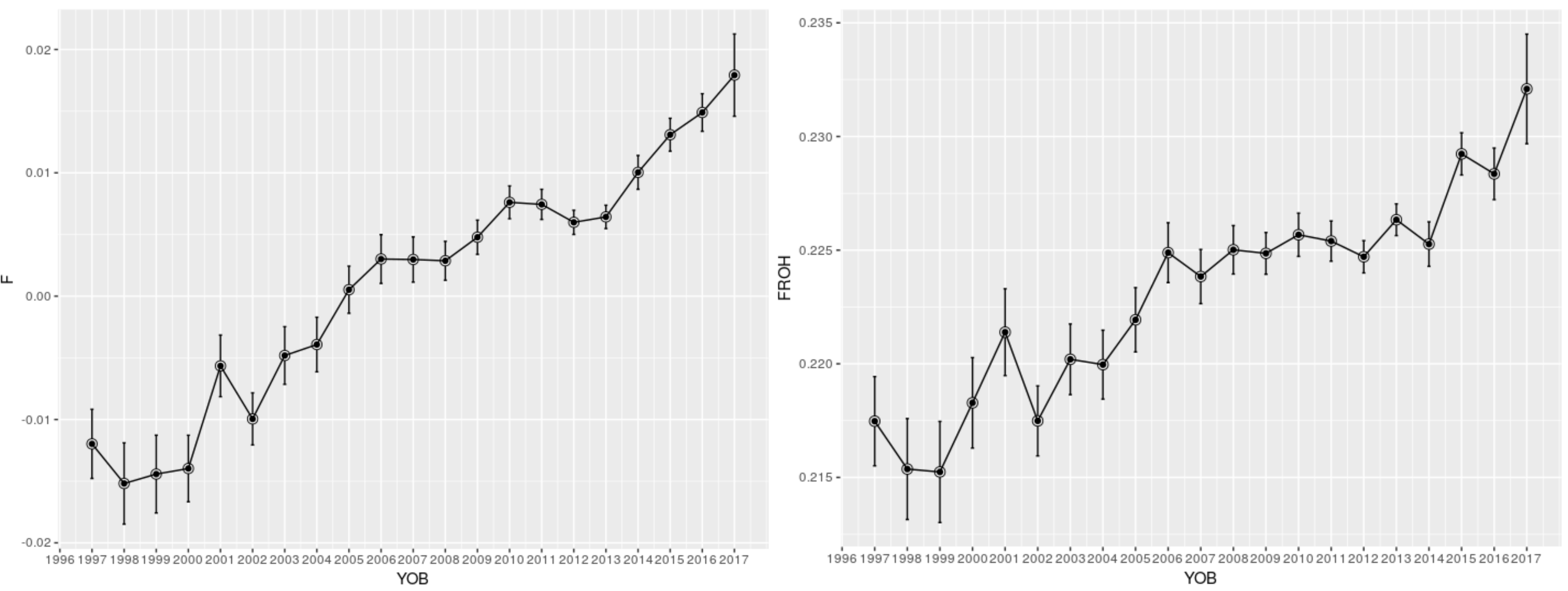

### S13 Figure
### Regional variation in inbreeding over time

Green – EUR
Blue – NAM
Red - ANZ

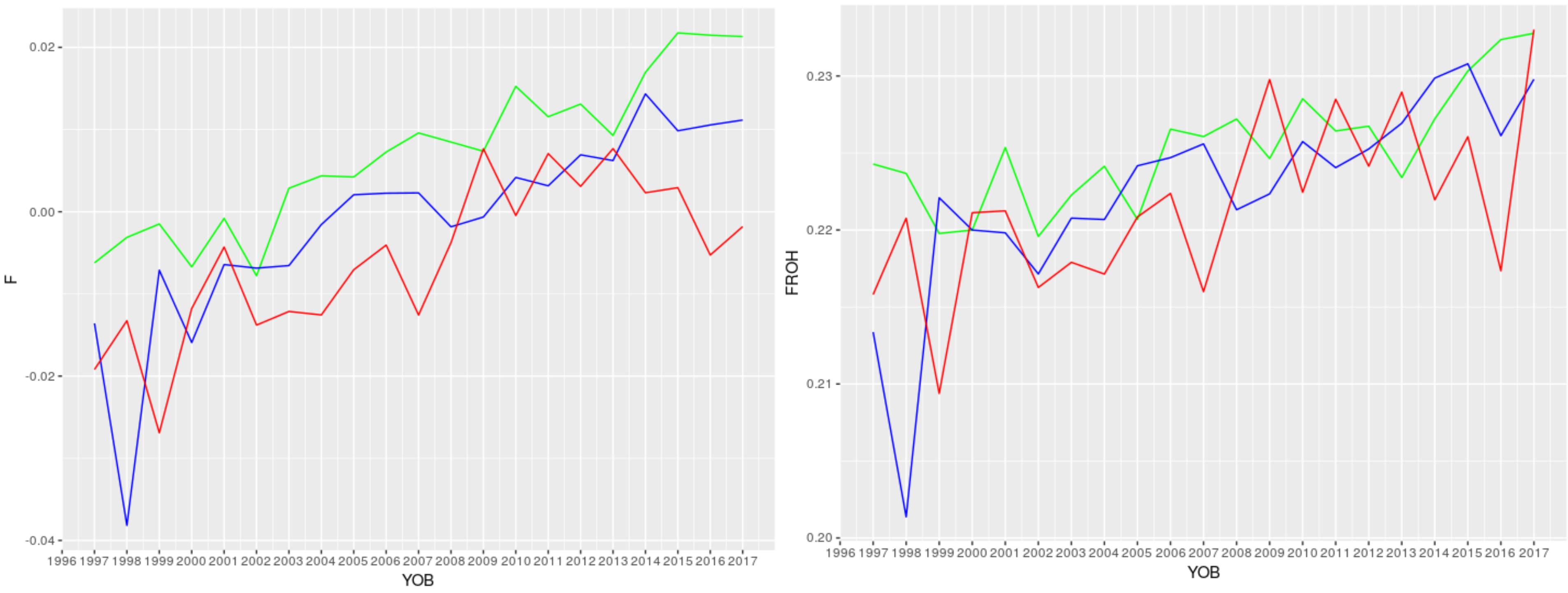

### S14 Figure
### Linear Regression fit for inbreeding by year of birth in Australasia (ANZ)

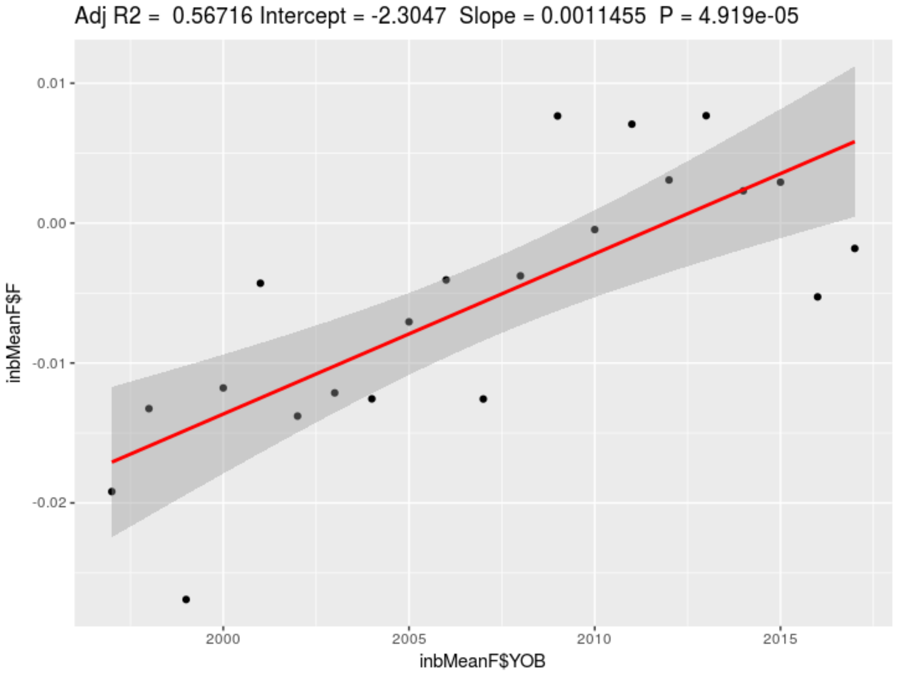


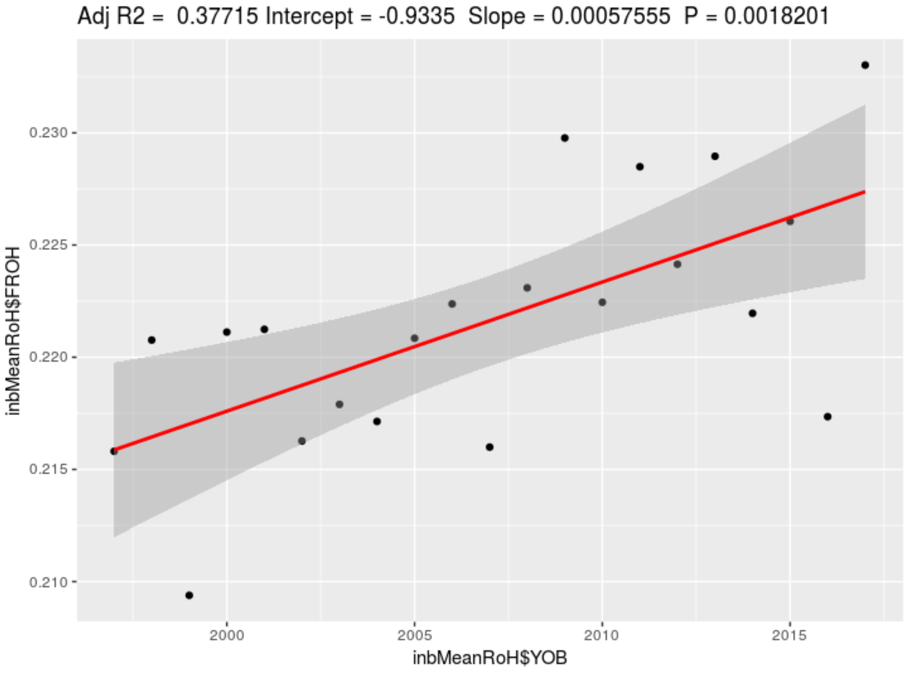

### S15 Figure
### Linear Regression fit for inbreeding by Year of birth in Europe (EUR)

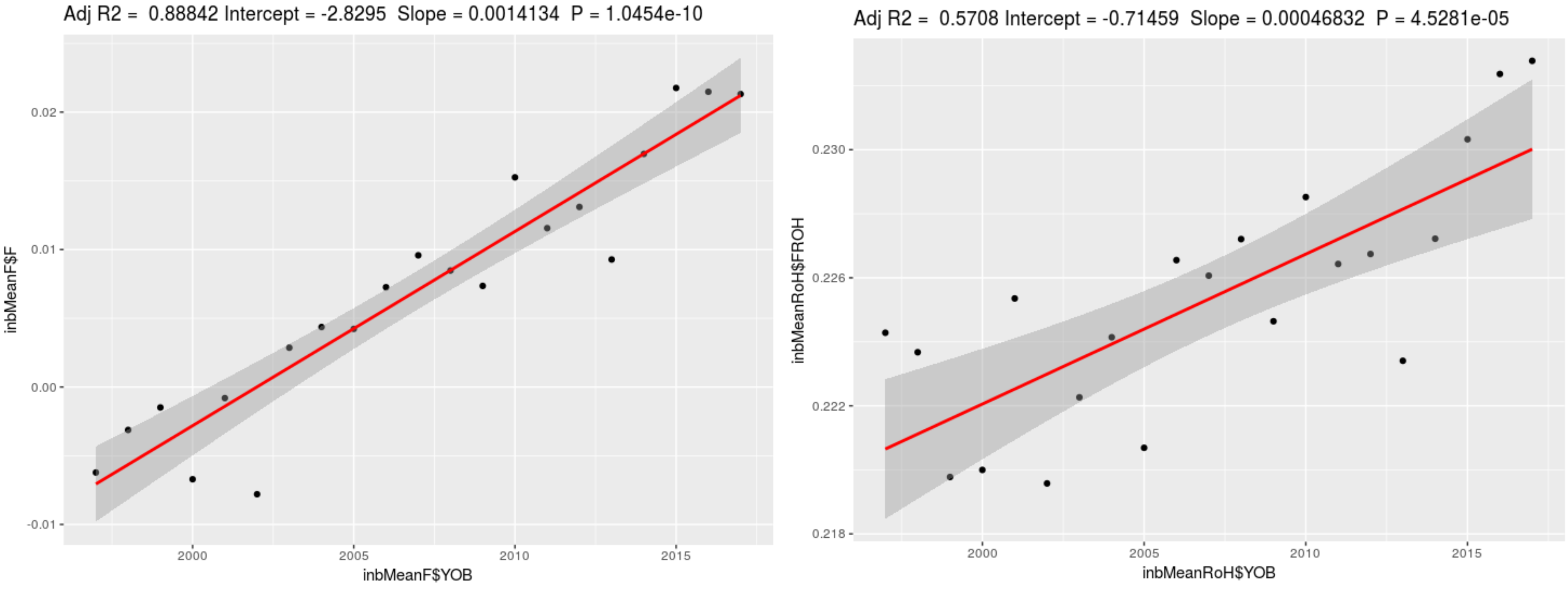

### S16 Figure

### Linear Regression fit for inbreeding by Year of birth in North America (NAM)

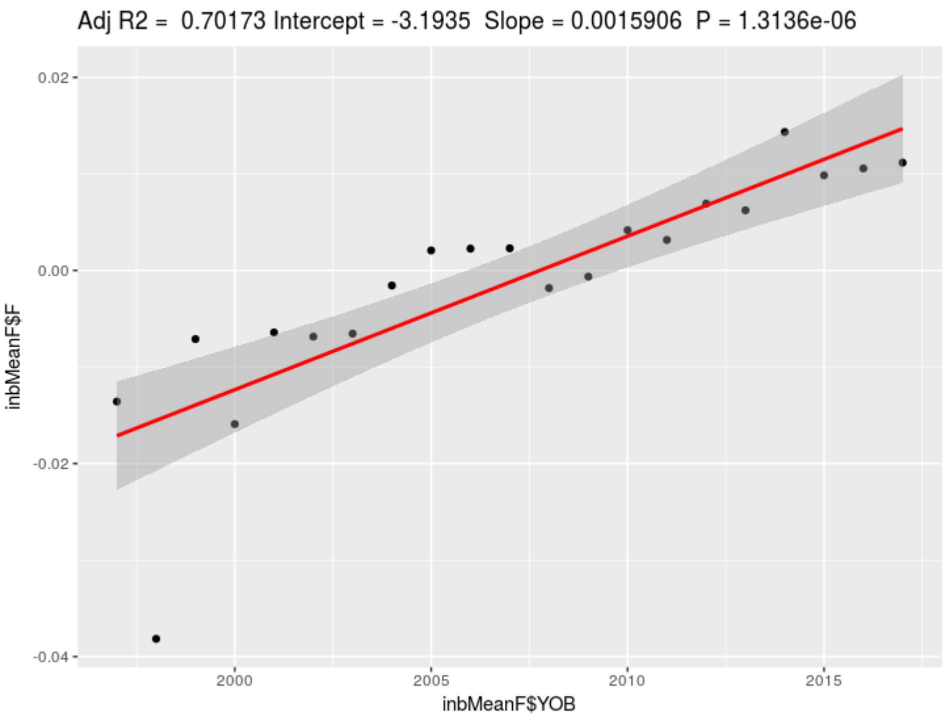


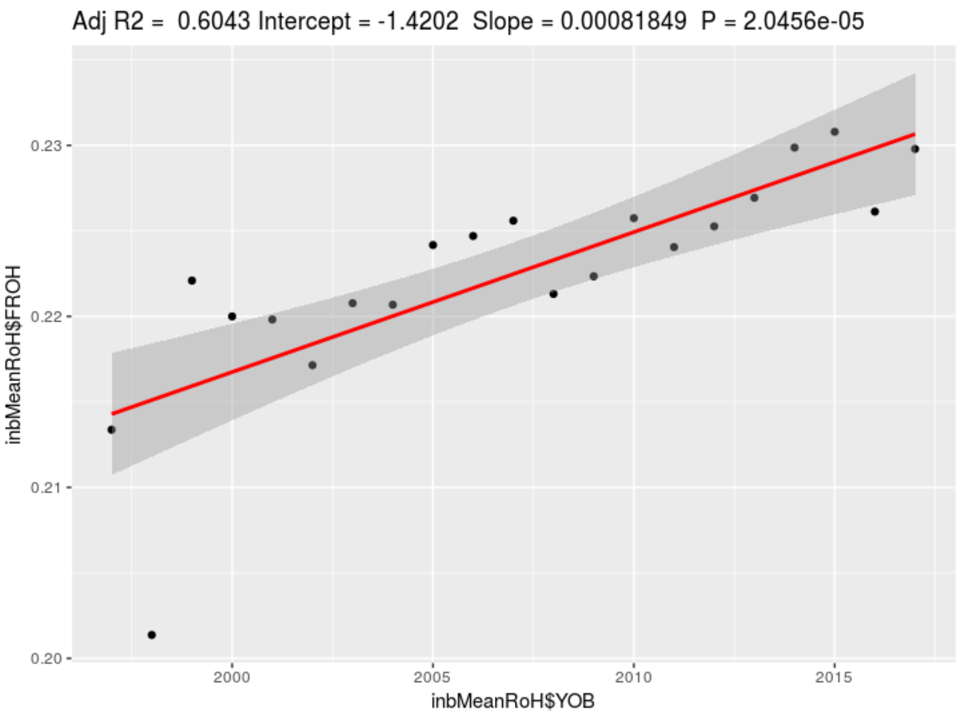

### S17 Figure
#### Manhattan plot for CSS and smoothed CSS values, showing significant gene regions under selection for Thoroughbreds

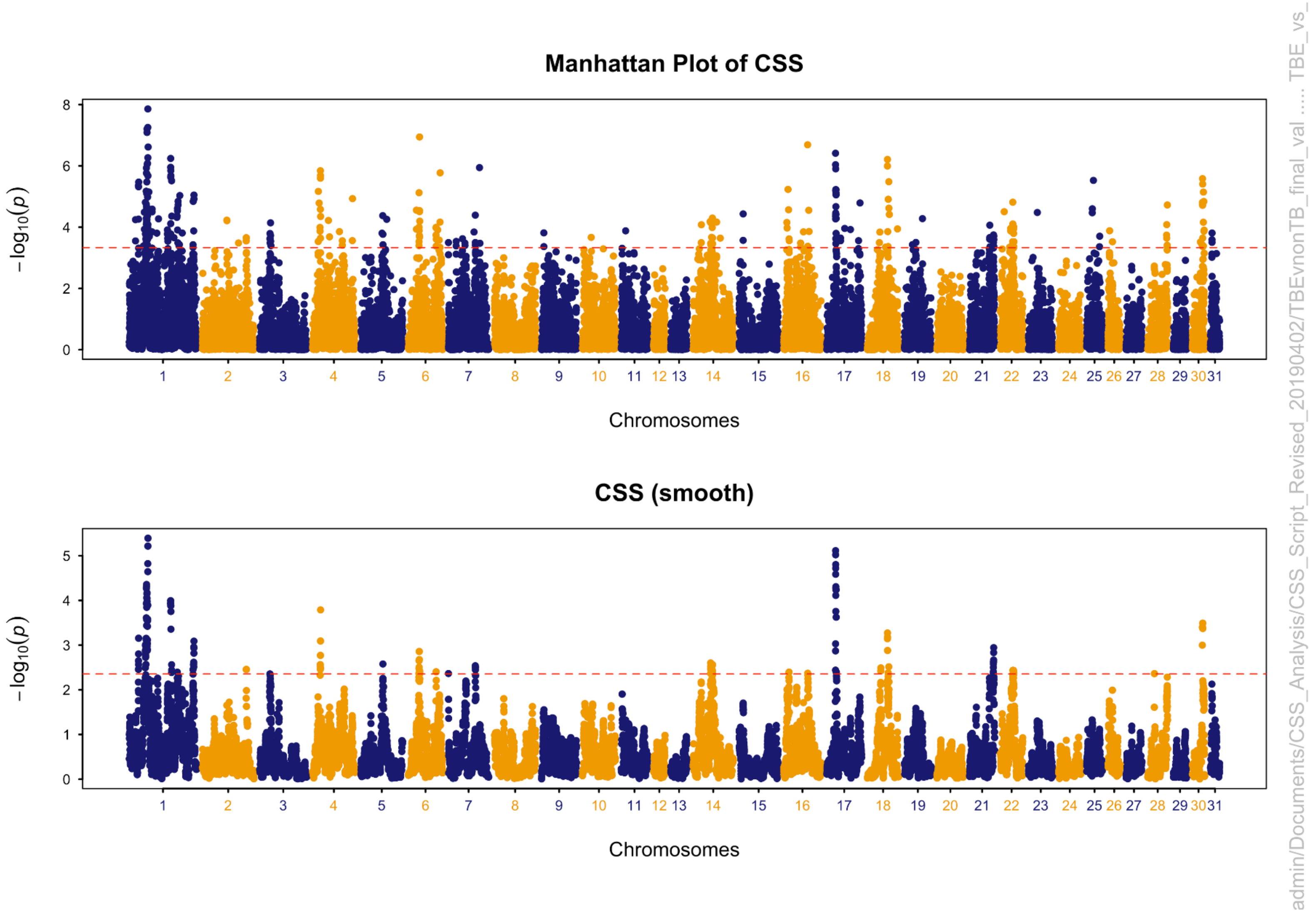

S18 Figure

Manhattan plot of the distribution of runs of homozygosity (Greater than 1Mb in length and SNP located within ROH in > 20% of the population) in the Thoroughbred population.

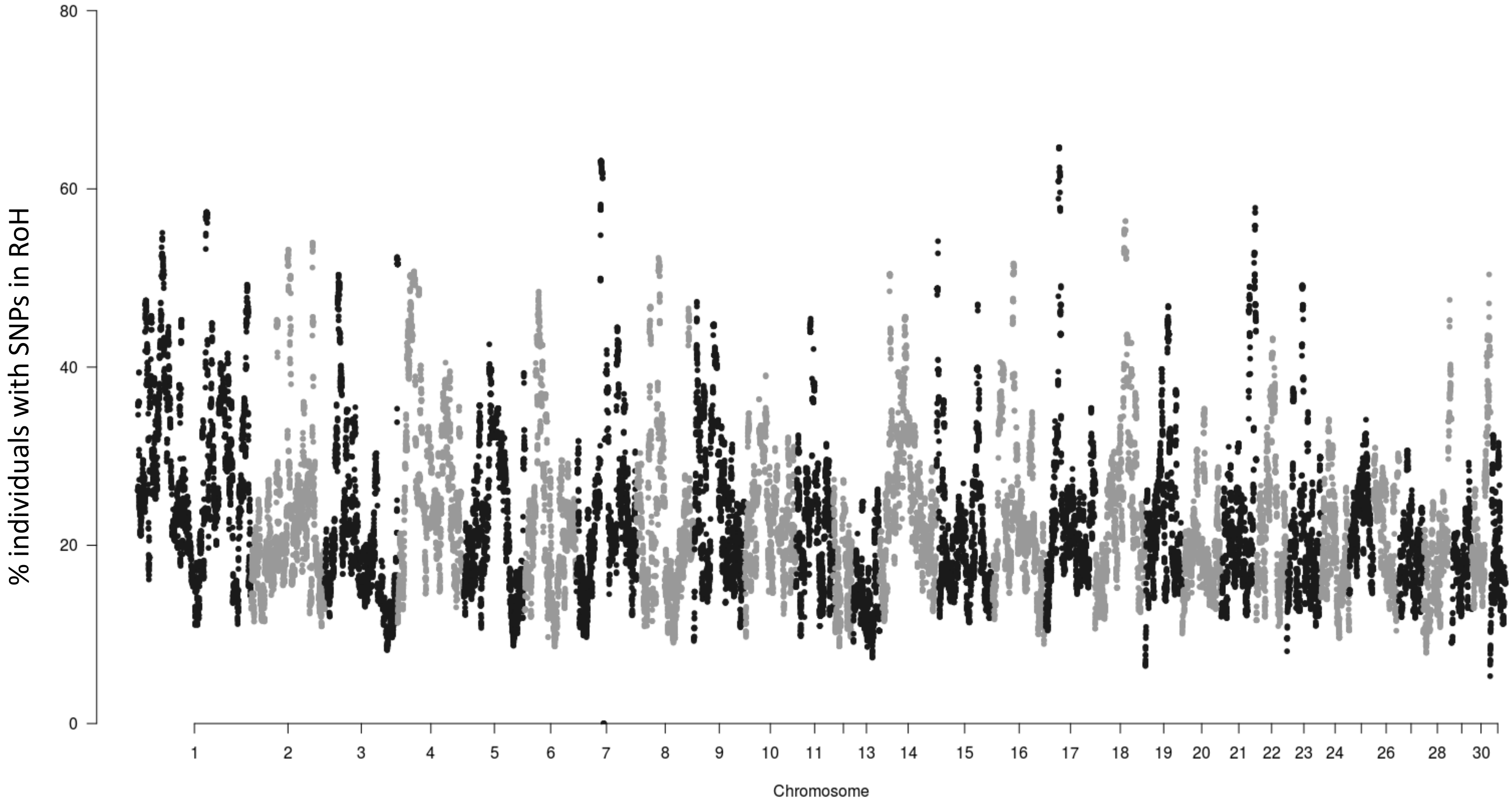

### S19 Figure
Comparison of pedigree-based estimates (x-axis) and the genomic estimate of relatedness (y-axis) of direct descendants of *Danehill*

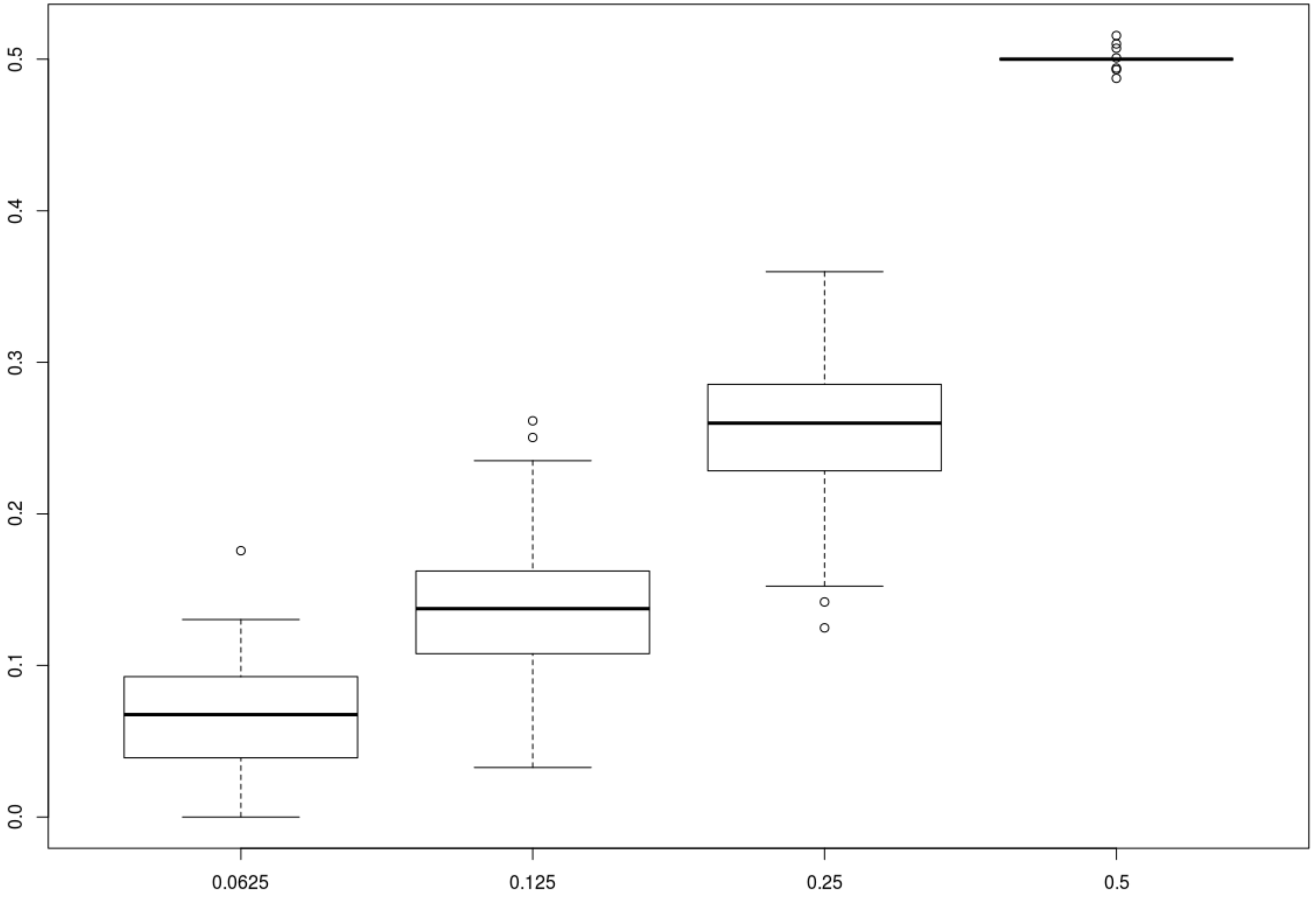
