## Supplemental_Text1 for "Genomic inbreeding trends in the global Thoroughbred horse population driven by influential sire lines and selection for exercise trait-related genes"

### S1 Text: Analysis of inbreeding and selection signatures in the Thoroughbred population

#### Stallion genotype reconstruction

Autosomal SNPs (46,478) were imputed for *n* = 127 sires based on genotyped progeny. Forty-three of these horses were also genotyped on at least one of the three SNP genotyping platforms. Concordance between inferred and SNP genotyping platform derived data ranged from 97.7-99.9% (S15 Table) with greater than 99% for 33 samples. Genetic distance was calculated using the pairwise identity by descent estimation function in Plink (1). Distance is defined as (IBS2 +0.5*IBS1)/(n SNP pairs).

#### Inbreeding in the Thoroughbred population

Annual mean inbreeding levels are provided in S16-17 Tables and S12 Figure. Regional variation in inbreeding over time is shown in S13 Figure. Results of linear models of the relationship between inbreeding and year of birth within each region are provided in S14-S16 Figures. The average inbreeding value within the modern breeding population (horses born since 2010 with offspring) is $F$ = 0.007 with no difference between stallions (*n* = 34) and mares (*n* = 75).

In addition to inferring inbreeding, analysis of runs of homozygosity (ROH) can identify genomic regions with the highest prevalence of ROH in the population potentially containing selected alleles. The top 1,000 SNPs ranked by the percentage of individuals with SNP located within ROH are provided in S2-S3 Tables and S18 Figure. However, ROH is not entirely informative on its own as alternate alleles of the same SNP may be identified in ROH in different animals. e.g. AA of the SNP BIEC2-5808 could be in a ROH in 20% of animals and GG in 26% of animals, appearing as ROH in 46% of animals. An additional limitation of ROH–based approaches lies in their underlying rule-based procedure. For instance the definition of the number of markers, segment length and proportion of allowed heterozygous markers is largely arbitrary and also dependent on SNP density. Nonetheless, evaluation of the top SNPs ranked by the percentage of individuals homozygous for the SNP within ROH >1Mb identified the neurotrimin (*NTM*) gene region in >63% of horses (S2 Table ). *NTM* has been highlighted as a hallmark of domestication in horses (2) and is associated with the chance of a Thoroughbred horse having a racecourse start (3), which may reflect its known function in neurodevelopment and the establishment of synapses (4).

**Pedigree analysis**

Within our dataset 97% of Thoroughbreds have *Northern Dancer* in their bloodline. This trend of rapid expansion of a dominant sire line is continuing. Within our dataset 410/743 horses born in ANZ between 2012 and 2017 can be traced back to *Danehill* (*Northern Dancer*'s grandson) within just three generations (i.e. 55% of Australian horses related through *Danehill* as a great-grandsire or closer). Within our dataset 1147/3298 horses born in EU between 2012 and 2017 can be traced back to *Sadler's Wells* (*Northern Dancer*'s son) within just three generations (i.e. 35% of EUR horses related through *Sadler's Wells* as a great-grandsire or closer). While pedigree data can be useful in highlighting broad trends in breeding practices multiple studies have shown that pedigree-based estimates of inbreeding and relatedness are less accurate than genomic methods (5-7). This disparity between pedigree and genomic relatedness can be seen among *Danehill* descendants - only those with *Danehill* appearing once in their pedigree were included, $n$ = 698 horses with year of birth (YOB) ranging from 1992 - 2017. A simplified pedigree-based estimation was used where relatedness to sire was estimated to be 0.5, relatedness to grandsire was estimated to be 0.25 and relatedness to great grandsire was estimated to be 0.125. Within the F1 generation (*Danehill* is sire; $n$ = 18) genomic relatedness calculated using the IBD function in Plink (1) is close to 0.5, in agreement with pedigree-based estimates. However, in subsequent generations there are substantial deviations from the pedigree estimate. Within the F2 generation (*Danehill* is grandsire; $n$ = 345) genomic relatedness ranged from 0.12 to 0.36 in contrast to pedigree-based estimates of 0.25, and 59% of horses from the F3 generation were as closely related to *Danehill* as those from the F2 generation (S19 Figure).

As a small number of sire lines are dominating Thoroughbred pedigrees there are also concerns regarding the loss of bloodlines with speculation that *The Byerley Turk* bloodline may soon be lost. Therefore it was of interest to look in detail at some of the remaining descendants of *The Byerely Turk* standing at stud today. We obtained blood samples from five stallions which trace back to *The Byerely Turk* on the male bloodline. These five stallions did not cluster separately on any of the PCA plots and inbreeding levels were comparable to those of other horses from the same regions and with the same year of birth. Recent analysis of Y chromosome variation {Felkel, 2019 #549} has established that there are likely more true *The Byerley Turk* lineages in the present population than previously thought; horses descended from *Galopin*, previously attributed to the *Darley Arabian* male lineage, in fact are descendants of *The Byerley Turk*.

#### ***CSS results***

Genome-wide distribution of the smoothed CSS ($-\log_{10}P$) for the comparison of the Elite Thoroughbred (TBE) *versus* Thoroughbred founder populations identified 15 significant candidate selected genomic regions (S1 Table, S17 Figure), defined as clusters of ≥5 SNPs among the top 1% of the smoothed CSS statistic result ($-\log_{10}P$). 462 genes were identified underlying these selection peaks and flanking regions (± 0.5 Mb) and 387 were used as input for IPA analysis. The top canonical pathway identified was Airway Inflammation in Asthma. A full list of pathways is provided in S12 Table.
